# Long read sequencing reveals a novel class of structural aberrations in cancers: identification and characterization of cancerous local amplifications

**DOI:** 10.1101/620047

**Authors:** Yoshitaka Sakamoto, Liu Xu, Masahide Seki, Toshiyuki T. Yokoyama, Masahiro Kasahara, Yukie Kashima, Akihiro Ohashi, Yoko Shimada, Noriko Motoi, Katsuya Tsuchihara, Susumu Kobayashi, Takashi Kohno, Yuichi Shiraishi, Ayako Suzuki, Yutaka Suzuki

**Affiliations:** Department of Computational Biology and Medical Sciences, Graduate School of Frontier Sciences, The University of Tokyo, Chiba, Japan; Division of Translational Informatics, Exploratory Oncology Research and Clinical Trial Center, National Cancer Center, Chiba, Japan; Division of Translational Genomics, Exploratory Oncology Research and Clinical Trial Center, National Cancer Center, Chiba, Japan; Division of Genome Biology, National Cancer Center Research Institute, Tokyo, Japan; Department of Pathology, National Cancer Center Hospital, Tokyo, Japan; Division of Cellular Signaling, National Cancer Center Research Institute, Tokyo, Japan

**Author notes:** To whom correspondence should be addressed. Yutaka Suzuki. Email address: Yoshitaka Sakamoto, Liu Xu, Masahide Seki, Toshiyuki T. Yokoyama, Masahiro Kasahara, Yukie Kashima, Akihiro Ohashi, Yoko Shimada, Noriko Motoi, Katsuya Tsuchihara, Susumu Kobayashi, Takashi Kohno, Yuichi Shiraishi, Ayako Suzuki, Yutaka Suzuki.

**Keywords:** Long read sequencing, Lung cancer, Local structural aberrations

## Abstract

Here we report identification of a new class of local structural aberrations in lung cancers. The whole-genome sequencing of cell lines using a long read sequencer, PromethION, demonstrated that typical cancerous mutations, such as point mutations, large deletions and gene fusions can be detected also on this platform. Unexpectedly, we revealed unique structural aberrations consisting of complex combinations of local duplications, inversions and micro deletions. We further analyzed and found that these mutations also occur *in vivo*, even in key cancer-related genes. These mutations may elucidate the molecular etiology of patients for whom causative cancerous events and therapeutic strategies remain elusive.

## INTRODUCTION

Recent cancer sequencing projects, such as the International Cancer Genome Consortium (ICGC) and The Cancer Genome Atlas (TCGA), have revealed causative mutations in various types of cancers^1,2^. Among them, lung adenocarcinomas are one of the most well-studied cancers regarding such “cancer driver” mutations^3–6^. In lung cancer patients, more than half of the cases have characteristic point mutations in the *EGFR* and *KRAS* genes or gene fusions in the *ALK, RET* and *ROS1* genes. These mutations are utilized as “biomarkers”, providing fundamental information about the most appropriate therapeutic strategy. Patients are separated based on their genomic mutation statuses and are matched to the most appropriate treatment^7–11^. Despite this general success, approximately 20-30 % of the lung adenocarcinoma patients remain undiagnosed with respect to their cancerous mutations^7^.

Current information on the cancer mutations has been mostly obtained by short read sequencing. Short read sequencing data, generally consisting of tens of millions reads of up to 200-300 bases in length^12^, are the most powerful in detecting point mutations such as single nucleotide variants (SNVs) and short indels^5,13^. Significant efforts have been made to enable the identification of fusion genes by short read sequences^14^. However, it is still difficult to detect more complex or larger-scale structural aberrations, such as chromosome aneuploidy, copy number aberrations and rearrangements solely based on short read sequencing data. There are inherent drawbacks even in the latest bioinformatics pipelines for this purpose, which hampers their practical use without careful validation^15^.

Recently developed long read sequencing technologies are changing this situation. Several pioneering papers have reported the precise analysis of complicated genomic regions, and large-range aberration detection is enabled by the long read sequencing. For example, a single molecule real-time (SMRT) sequencer, PacBio RS, has been utilized to analyze *BCR-ABL1* rearranged transcripts and their TKI-resistant mutations in chronic myeloid leukemia (CML)^16,17^. In Chromophobe Renal Cell Carcinoma (ChRCC), structural alterations in the *TERT* promoter region have been identified and characterized by PacBio sequencing^18^. Very recently, a nanopore-type sequencer, MinION, was first utilized to characterize the pathogenic sequence expansion of intronic repeats in Benign Adult Familial Myoclonic Epilepsy (BAFME)^19^. Particularly for cancer applications, we and others have shown that cancer-associated structural variants (SVs) could be detected by nanopore sequencing approaches^20,21^. Additionally, transcriptome sequencing by MinION has been shown to provide a powerful analytical platform, where the complete splicing pattern of a given mRNA can be thoroughly represented by a single read^22^. Further improvements in MinION sequencing have been achieved by the parallelization of the nanopores in a given flow cell, a platform named PromethION. PromethION can now produce over 100 Gb reads per flow cell.

In this study, we attempted long read sequencing of whole human cancer genomes using PromethION. We first demonstrate that PromethION sequencing can identify point mutations as well as large structural aberrations and fusion genes relatively easily. Moreover, we unexpectedly identified that mutations containing complex combinations of small and middle-sized structural aberrations are quite common, constituting a previously undefined unique class of mutations. Hereafter, we will call those mutations Cancerous Local Copy-number Lesions (CLCLs). These CLCLs resided even within the key cancer genes or drug target genes, such as the *STK11, NF1 SMARCA4* and *PTEN* genes. Additionally, taking advantage of long read sequencing, we characterized the full-length transcript structures by full-length cDNA sequencing of the transcriptome.

We initially used lung cancer cell lines for which we had previously collected detailed information on multi-omics features, such as whole-genome sequencing, RNA-seq, and ChIP-seq of Illumina reads^23^. Then, we used clinical samples to demonstrate that those CLCLs are not restricted to cell lines.

## RESULTS

### Long read sequencing of cancer cell lines

We conducted long read and whole-genome sequencing analysis using the nanopore type sequencers, MinION and its latest high-throughput derivative, PromethION. We first validated the performance of the new PromethION instrument by sequencing the genome of LC2/ad, which is a lung cancer cell line derived from a Japanese lung adenocarcinoma patient^24,25^ (**Fig. 1** and **Supplementary Table S1**). As a reference, we collected the whole-genome sequencing data from a total of 33 MinION runs (R9.5 flow cells) to cover the whole human genome at an overall sequencing depth of 31× by a total of 7,282,846 reads (93,813,338,154 base pairs (bp)). The maximum length and N50 length of the reads were 2,495,160 bp and 30,606 bp, respectively. In total, 67.5 % of the reads were mapped to the human reference genome UCSC hg38 using Minimap2^26^. The calculated overall sequence identity was 82 % on average. The average length of the mapped reads was 16,452 bp, which was significantly longer than previous long read whole human cancer genome sequencing analyses^27–29^ (**Table 1**). The PromethION sequencing required approximately three flow cells to generate a total of 10,064,668 reads (100,440,433,160 bp) for an overall coverage of 33× (**Fig. 1**) (this number decreased for subsequent cell types; see **Methods** for more details). The maximum length and N50 length of reads were 987,834 bp and 32,710 bp, respectively. Using Minimap2, 69.4 % of the reads were mapped to the reference genome. The average length of the mapped reads was 13,620 bp, and the average identity was 85 % (**Table 1**). Notably, because sample preparation need not be performed for each run, the required total amount of starting DNA used for PromethION could be reduced by more than tenfold compared with MinION.

**Table 1.**
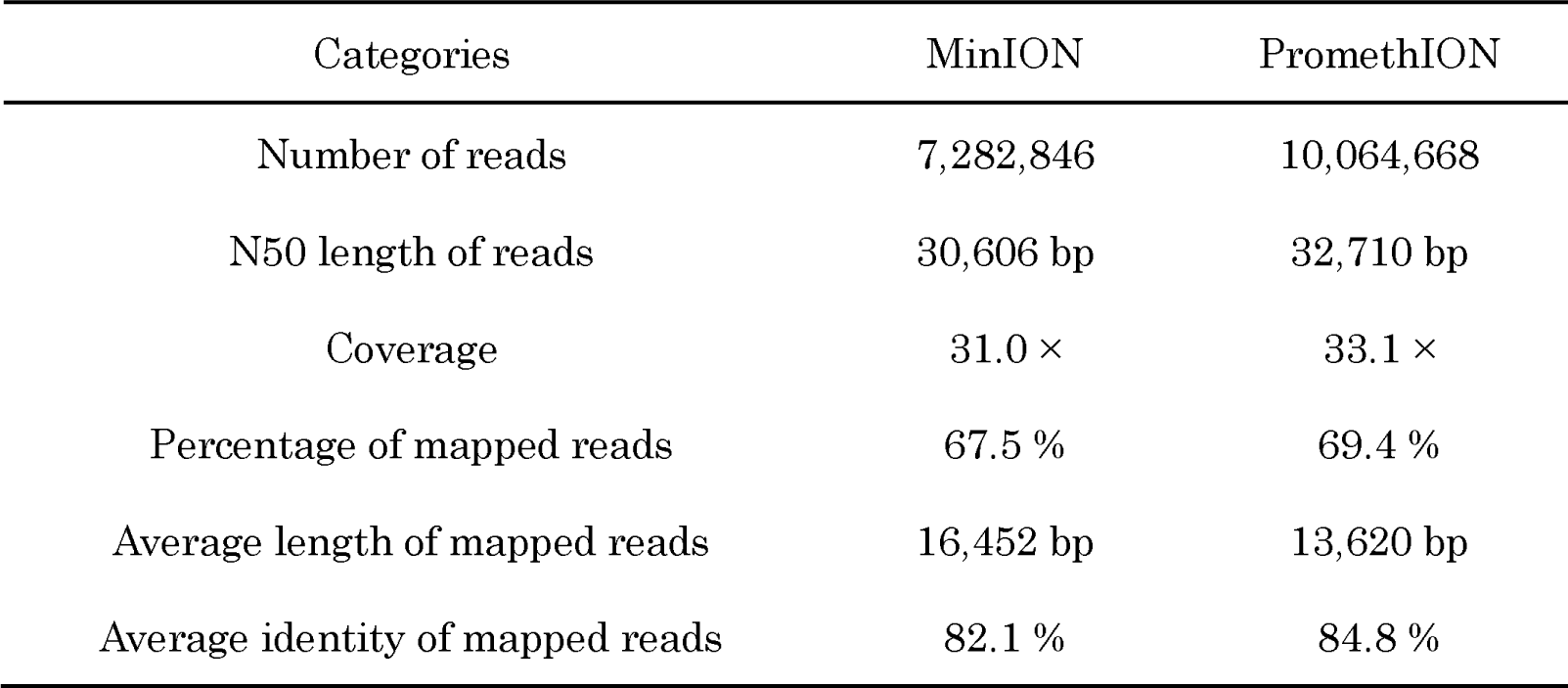
General statistics of nanopore sequencing in LC2/ad.

**Figure 1.**
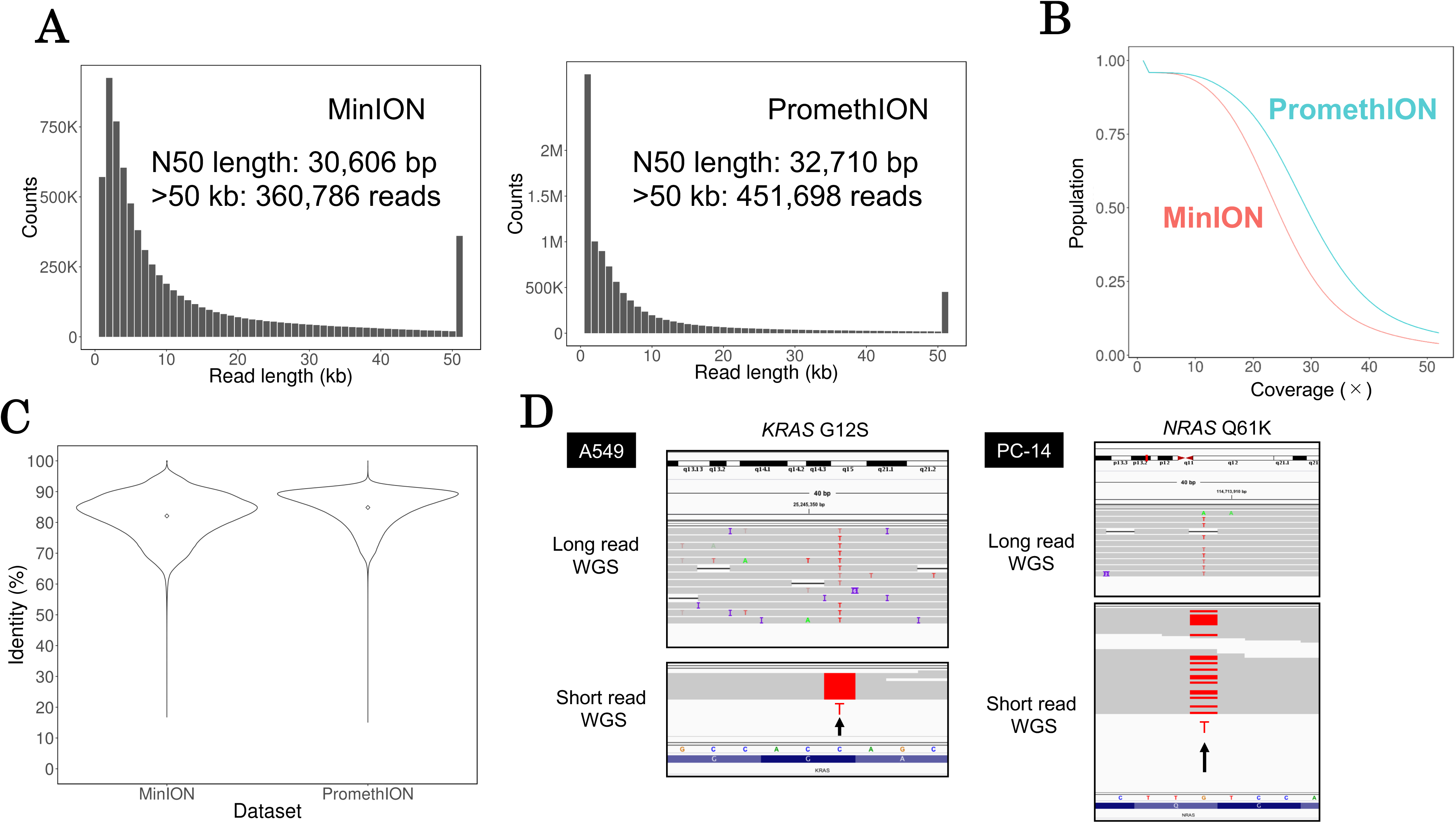
Long read sequencing of cancer genomes. (**A**) Left: raw read length distribution of LC2/ad MinION sequencing. Right: raw read length distribution of LC/2ad PromethION sequencing. Both MinION and PromethION datasets have many long reads (for example, over 50 kb, 361 K reads and 452 K reads, respectively). (**B**) Cumulative depth curve of LC2/ad. Blue: PromethION, Red: MinION. More than 50 % of the human genome region was covered by more than 20× in sequencing depth in both of the datasets. (**C**) Violin plots of identity of LC/2ad MinION sequencing (left) and PromethION sequencing (right). The points in the violin plots indicate the average identities of the MinION and PromethION data (82.1 %, 84.8 %, respectively, **Table 1**). The identities were concentrated at more than 80 % in both datasets. (**D**) IGV image of point mutations in the driver genes *KRAS* and *NRAS* in MinION/PromethION sequencing (upper) and in Illumina sequencing (lower). *KRAS* point mutation (G12S) is a driver mutation in A549 cells and *NRAS* point mutation (Q61K) is a driver mutation in PC-14 cells.

To examine whether PromethION sequencing was compatible with MinION sequencing, we compared the features of the obtained two datasets. The overall distribution of read lengths was similar (**Fig. 1A**). Both datasets included a substantial fraction of long reads over 50 kb (MinION: 360,786 reads, PromethION: 451,698 reads). Detailed analysis of the mapping results showed that more than 50 % of the human genome region was covered at more than 20× sequencing depth in both of the datasets (**Fig. 1B**). For the sequence accuracy, both datasets showed an overall fidelity of more than 80 % (**Fig. 1C**), which is similar to that of a previous study^30^. We concluded that PromethION should be an effective analytical method for whole cancer genome sequencing.

Having finished the initial evaluation of the data obtained from MinION and PromethION, we scaled the MinION and PromethION sequencing for an additional four lung cancer cell lines (A549, RERF-LC-KJ, RERF-LC-MS and PC-14; detailed cellular profiles are described **Supplementary Table S1**). The data production proceeded similarly to the case of LC2/ad cells, for example, in the sequencing of RERF-LC-KJ, 5,986,875 reads were generated (57,062,227,853 bp, at 18.5×), with the max and N50 lengths of the reads being 922,768 bp and 23,442 bp, respectively. Other detailed statistics are shown in **Supplementary Figure S1** and **Supplementary Table S2**.

To evaluate the quality of the sequence data at the individual base level, we examined the known driver mutations of the corresponding cells by manually reviewing the mapping results with Integrative Genomics Viewer (IGV)^31,32^. In A549, eleven reads illustrated the cancerous mutation *KRAS* G12S as the point mutation (left, **Fig. 1D**). In PC-14, eight reads represented the driver *NRAS* Q61K point mutation (right, **Fig. 1D**). Conversely, we also confirmed the absence of any driver mutations in the RERF-LC-KJ and RERF-LC-MS cell lines at well-known driver genes. All of these results are consistent with those of previous reports^23^. These results collectively indicated that mutation calling at the single-base level is also possible using only the long read sequencer, at least when the cancer cell contents are as high as in the cultured cells.

### Identification of large-scale genomic aberrations

Using the long read sequencing data, we then attempted to detect structural aberrations larger than point mutations (**Fig. 2A**). From the MinION/PromethION dataset of LC2/ad, we successfully identified 12 reads directly overlapping the junction point of the *CCDC6-RET* fusion gene, which is the known “cancer driver mutation” for this cell line^24,25^ (**Fig. 2B**; for details of the bioinformatics pipeline, see **Methods**). We further attempted to identify large deletions. A large deletion around the *CDKN2A* gene, which is a well-known tumor suppressor gene^3^, was previously reported to occur in LC2/ad, A549 and PC-14 cells^23^ (**Fig. 2C**). Using the MinION/PromethION datasets in this study, we re-confirmed the deletion of this gene in the respective cells. In addition, we found that the precise junction point of each of the *CDKN2A* deletions was different between the cell types. Large deletions in other cancer-related genes are described in **Supplementary Figure S2**.

**Figure 2.**
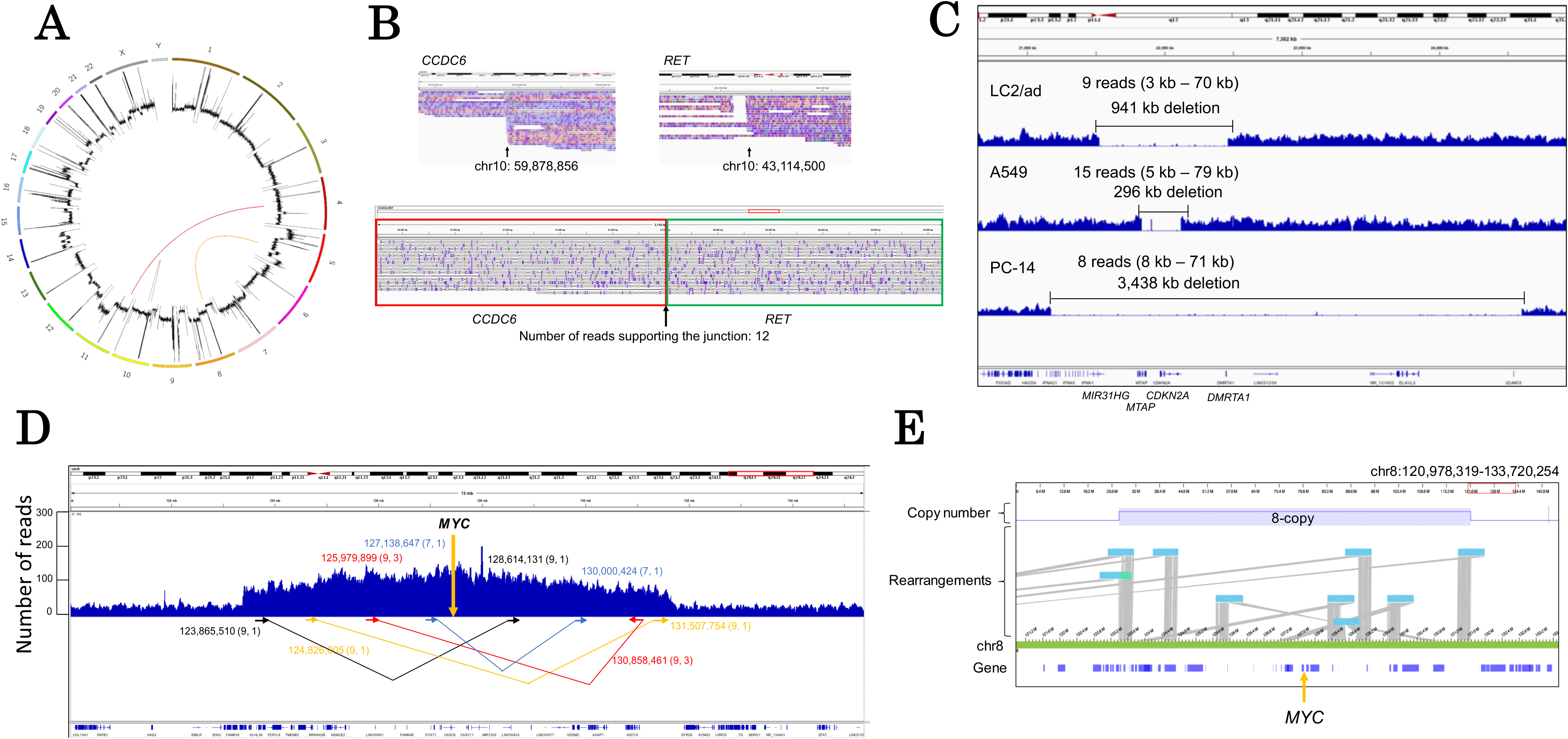
Long read sequencing reveals structural variants. (**A**) Circos plot of novel rearrangement candidates in LC2/ad. The second layer of the Circos plot indicates the sequencing depth by MinION. Detailed information on this rearrangement is shown in **Supplementary Table S3**. (**B**) IGV image of the *CCDC6-RET* fusion gene of LC2/ad MinION sequencing. The *CCDC6-RET* fusion gene is a driver mutation of LC2/ad cells. The number of reads supporting the junction is twelve. (**C**) IGV image of a large deletion including *CDKN2A* of LC2/ad, A549, and PC-14. The deletion of LC2/ad spanned approximately 941 kb. The deletion of A549 spanned approximately 296 kb. The deletion of PC-14 spanned approximately 3,438 kb. The parentheses indicate the range of length of reads supporting the deletions. (**D**) Depth plotting around the *MYC* gene of LC2/ad. Arrows indicate junction candidates of the amplification supported by MinION and Illumina reads. (): the number of supporting MinION reads (left) and Illumina paired-end reads (right). (**E**) The 8-copy *MYC* region of LC2/ad represented by the optical mapping method. Optical maps with rearrangements and chromosome 8 (reference) are represented in light blue and yellow-green, respectively. The orange arrow indicates the *MYC* gene.

We could also detect novel gene fusions by employing the split alignment method (see **Methods**). We identified three novel rearrangements, which were further validated by the Illumina short reads (**Fig. 2A** and **Supplementary Fig. S3**). These genes were fused to *NELL1-CCSER1* and *EFNA5-IKBKB* in LC2/ad and *UTS2B-GRM4* in RERF-LC-KJ. In each of these cases, the long read sequencing precisely identified the junction at single-base resolution (**Supplementary Table S3**).

We further attempted to decipher perhaps the most difficult case, the rearrangement of the *MYC* gene. We identified copy number aberrations of the *MYC* gene in LC2/ad^23^. The amplification was estimated to extend over approximately an 8 Mb locus having the *MYC* gene at the center. Even using long read sequencing, it was still difficult to completely reconstruct its structure, which included complex rearranged patterns, expanding to 8 Mb in chromosome 8 at an estimated aneuploidy of eight (**Fig. 2D**). Particularly for the *MYC* region, we attempted to identify the correct structure by the optical mapping method, Bionano Saphyr. Even using the Saphyr, the precise structure of the *MYC* region remained elusive, though the results from this analysis support the *MYC* amplification spanning the 8 Mb region with approximately 8 copies (**Fig. 2E**).

### Identification of a new class of cancerous local genomic lesions, CLCL

During the attempts to identify the above structural aberrations of the established classes, we unexpectedly found a new type of local structural aberration (**Fig. 3**). These aberrations consisted of complex combinations of copy number changes, inversions and deletions. As it appears that these aberrations do not precisely belong to the above categories, we named them Cancerous Local Copy-number Lesions (CLCLs). As we will describe below, we found it difficult to identify and characterize these CLCLs regarding their precise junctions solely based on short read sequencing, even though some suggestive data could be occasionally obtained.

**Figure 3.**
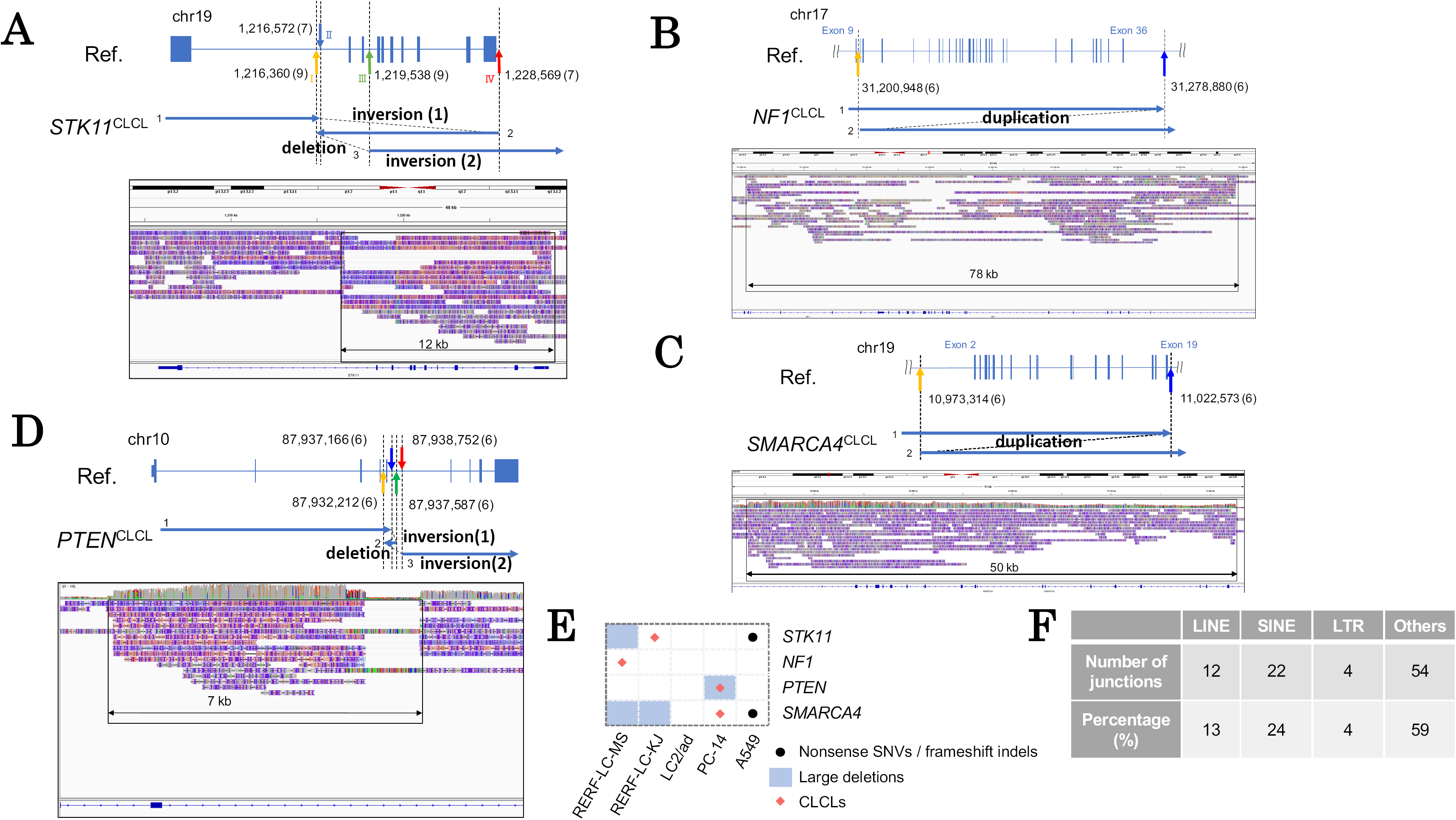
Identification and characterization of CLCL. (**A**) Structure of the *STK11* CLCL in RERF-LC-KJ. The *STK11* CLCL was constructed by a combination of local inversions. We can trace the CLCL structure following the ordered arrows, and the junctions are indicated by colored arrows. The CLCL spanned 12 kb in the human reference genome. (**B**) Structure of the *NF1* CLCL in RERF-LC-MS. The structure of the CLCL was a tandem duplication between the junctions (indicated by a yellow arrow and a blue arrow). The CLCL spanned 78 kb in the reference genome. (**C**) Structure of the *SMARCA4* CLCL in PC-14. The structure of the CLCL was a tandem duplication of the junctions (indicated by a yellow arrow and a blue arrow). The CLCL spanned 50 kb in the reference genome. (**D**) Structure of the *PTEN* CLCL in PC-14. The structure of the CLCL was a combination of a local inversion and deletion. We can trace the CLCL structure following the ordered arrows, and the junctions are indicated by colored arrows. The CLCL spanned 7 kb in the reference genome. (**E**) Summary of mutation types of four-cancer-related genes in five cell lines. (**F**) The number of CLCL junctions in each category of genomic contexts.

The first example was found in the *STK11* gene locus. In our previous study of lung cancer whole-genome sequencing using Illumina, we noticed a possible local copy-number lesion in the *STK11* gene region in RERF-LC-KJ cells. The sequencing depth increased from the middle of intron 1 to the end of the gene^23^. There were short read split tags (see Methods for details), suggesting that the inversions may occur in this region. Despite the substantial number of sequencing reads mapped in this region, we could not reconstruct its precise structure.

We examined the long reads to decipher the aberration in the *STK11* gene locus (**Fig. 3A**). It revealed the aberration as follows: The first rearrangement occurred as an inversion starting from intron 1 (chr19: 1,216,572; breakpoint II) and jumping downstream of the gene (chr19: 1,228,569; breakpoint IV). The inverted sequence continued back to the middle of the intron 1 (chr19: 1,216,360; breakpoint I), which was 212 bases upstream of the initial breakpoint II. Then, the sequence reverted back and jumped to intron 3 (chr19: 1,219,538; breakpoint III). The following sequence continued to the end of the gene locus. The detected junctions, breakpoints II/IV and I/III, were represented by seven and nine PromethION reads, respectively. When we re-examined the Illumina reads, the sequencing depth increased at the two regions, between breakpoints I and II and between breakpoints III and IV (boxed region in **Fig. 3A**). We also looked for the short reads using the soft-clipped method. We found that it was difficult to detect two of the breakpoints, I and III, using the short read split tags, partly because the junctions were resided in the repetitive regions.

### Identification of CLCLs in other genes and cell lines

To more generally identify CLCLs in other loci in all lung cancer cell lines, we constructed a new analytical bioinformatics pipeline (see **Supplementary Fig. S4** and **Methods**). Briefly, we utilized the information of the split alignments from the mapping results. We sorted the mapping information by the position of the reads and extracted the CLCL candidates. The associated reads were reassembled to reconstruct their structures.

As a result, we successfully identified the following numbers of CLCLs in the other cell lines as well: sixteen in LC2/ad, one in A549, seven in RERF-LC-KJ, seven in RERF-LC-MS, and eleven in PC-14 (**Table 2**). Importantly, CLCLs were found to occur even in key cancer genes, such as the *STK11, NF1, SMARCA4* and *PTEN* genes. The aberrant structures varied, and most of them would not be easily detected by the conventional short-read-based approaches because of their complex structures and the size of the affected regions. A relatively simple one was that which was detected in the *NF1* gene in RERF-LC-MS cells (**Fig. 3B**). This was a tandem duplication of the region between intron 9 (chr17:31,200,948) and the downstream region of the last exon 36 (chr17: 31,278,880; it was supported by six reads at the junction). In another case, the structure of the *SMARCA4* CLCL showed a duplication from intron 1 (chr19:10,973,314) to intron 20 (chr19: 11,022,573; supported by eight reads at the junction; **Fig. 3C**). A more complex case was found in the structure of *PTEN* in PC-14. This CLCL was found to be a combination of inversion and deletion (**Fig. 3D**). In these relatively simple cases, remapping of the Illumina short reads to the discovered junctions validated the precise identification of the reconstructed structure.

**Table 2.**
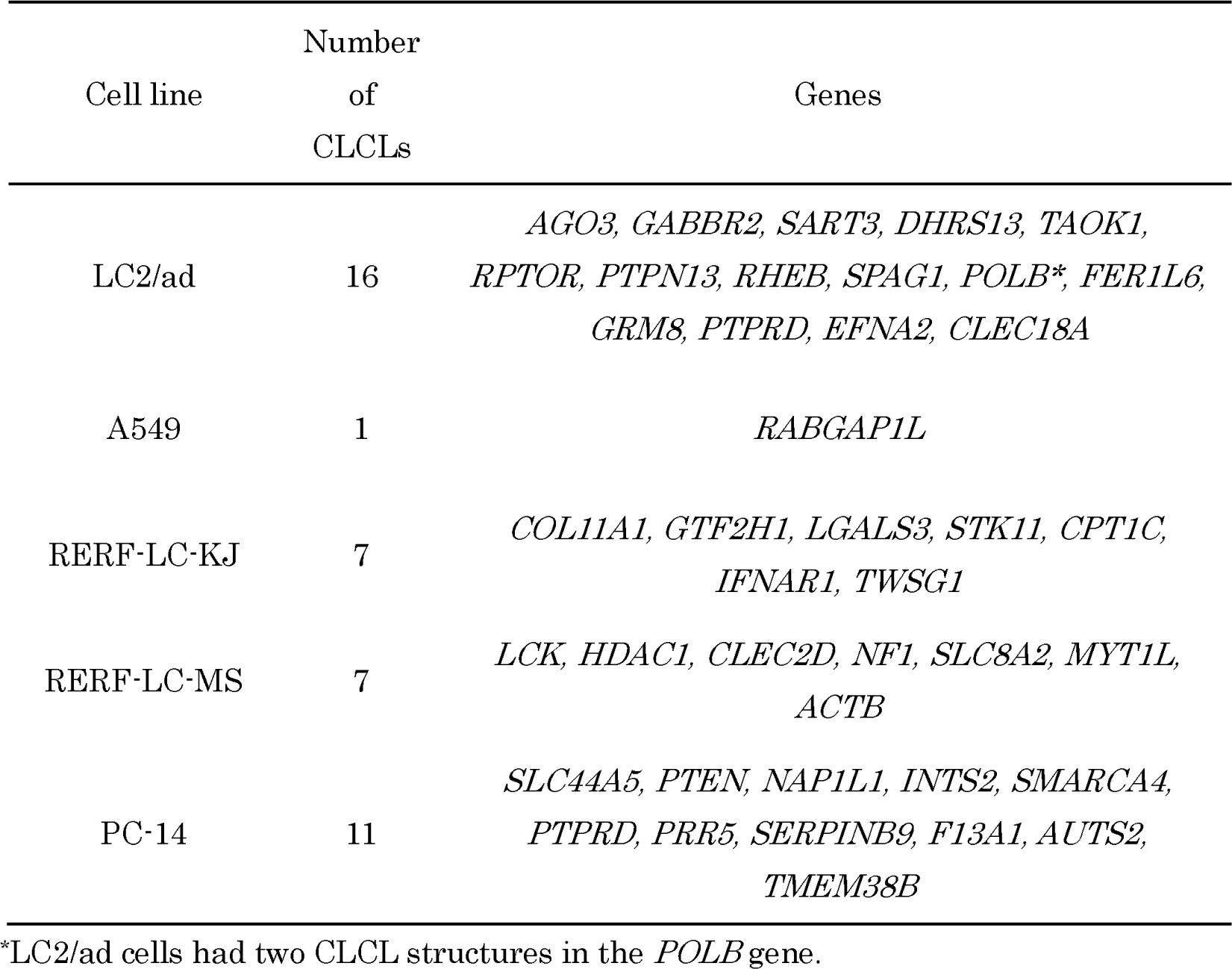
Summary of CLCLs detected in lung cancer cell lines.

Indeed, although the presence of these mutations was partly suspected in a previous study^23^, their precise structures remained elusive before this study. We and others had previously suspected the presence of large deletions, frameshift indels and splice site mutations based on short read sequencing for those cases. However, by conventional aberration detection based on the short reads, we could not detect some cases, which were first identified as CLCLs in this study (indicated by black dots in **Fig. 3E**).

We also examined the genomic context of the CLCLs. In total, 64 % (28/44) of the CLCLs had at least one junction overlapping with a long interspersed nuclear element (LINE), short interspersed nuclear element (SINE) or long terminal repeat (LTR), and 13 %, 24 % and 4 % (12/92, 22/92, and 4/92, respectively) of the junctions of the CLCLs were in a LINE, SINE or LTR, respectively (**Fig. 3F**). It is possible that their unique locations may hamper the precise identification of CLCLs by short read sequencing.

### Aberrant transcriptional events associated with CLCLs

After the new CLCL-type aberrations were identified in a number of key genes in a number of cell types, the immediately raised question was in what manner they have transcriptional or epigenomic consequences.

To characterize how the CLCL aberrations are reflected in the transcriptomes, we newly generated and analyzed full-length cDNA sequencing data using MinION. We also utilized the previous Illumina short read RNA-seq and ChIP-seq data. In RERF-LC-KJ cells, short read sequences indicated that the *STK11* transcript is abnormally spliced at intron 1 and that transcription jumped just before the CLCL structure^23^. MinION reads representing the full-length transcripts further specified the precise splice pattern and the transcription termination sites (**Fig. 4A**). For almost all of the transcripts, the first splicing occurred at the abnormal position (from chr19:1,216,268) and transcription occurred according to the CLCL structure (RNA-seq reads covered breakpoints II-IV from chr19:1,216,572 to chr19:1,228,569). Some aberrant transcription was also observed within the downstream CLCL region (middle panel, **Fig. 4A**). Such an aberrant transcription pattern was not observed in PC-14 cells, where the *STK11* gene is wild-type (lower panel, **Fig. 4A**).

**Figure 4.**
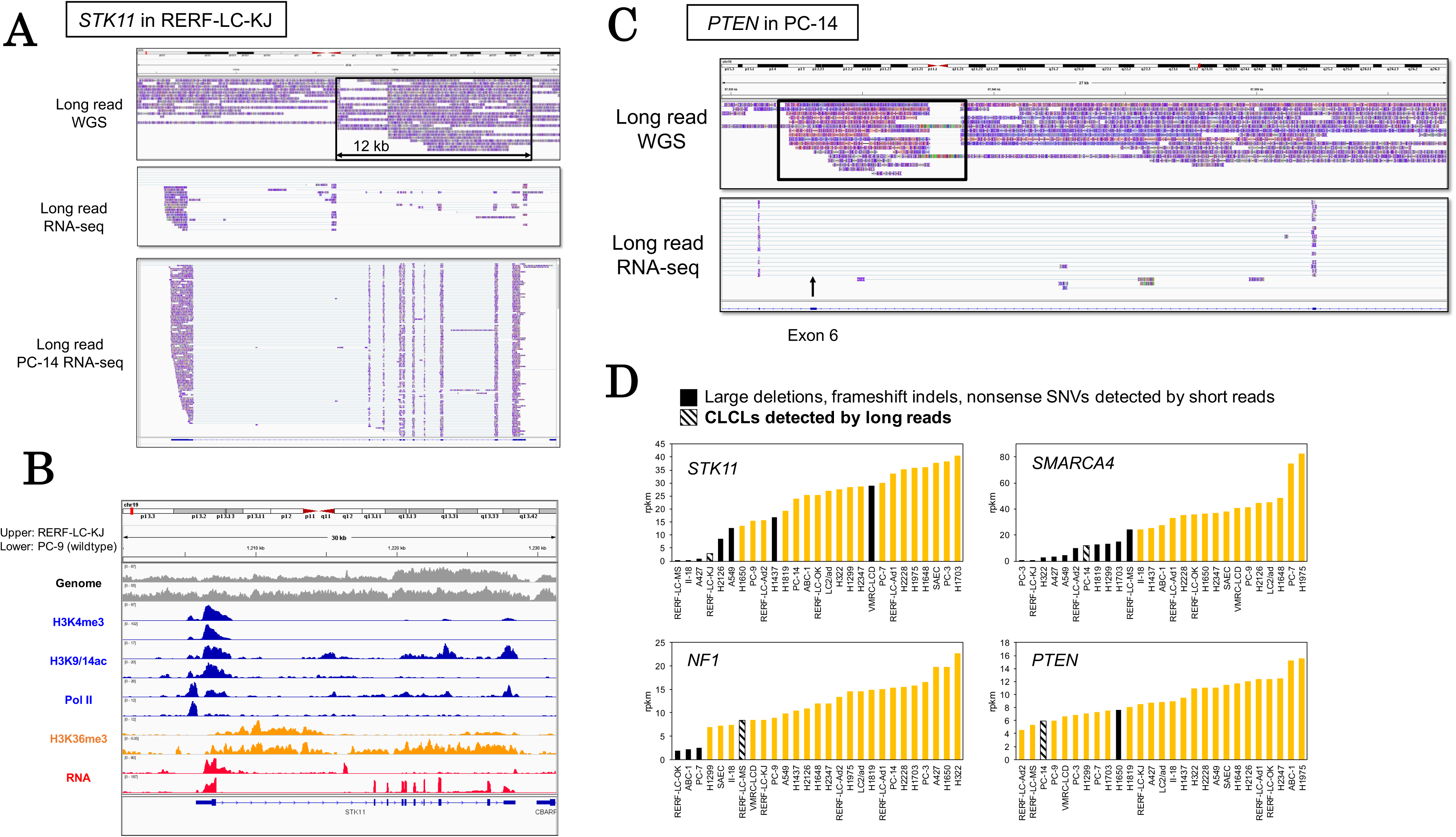
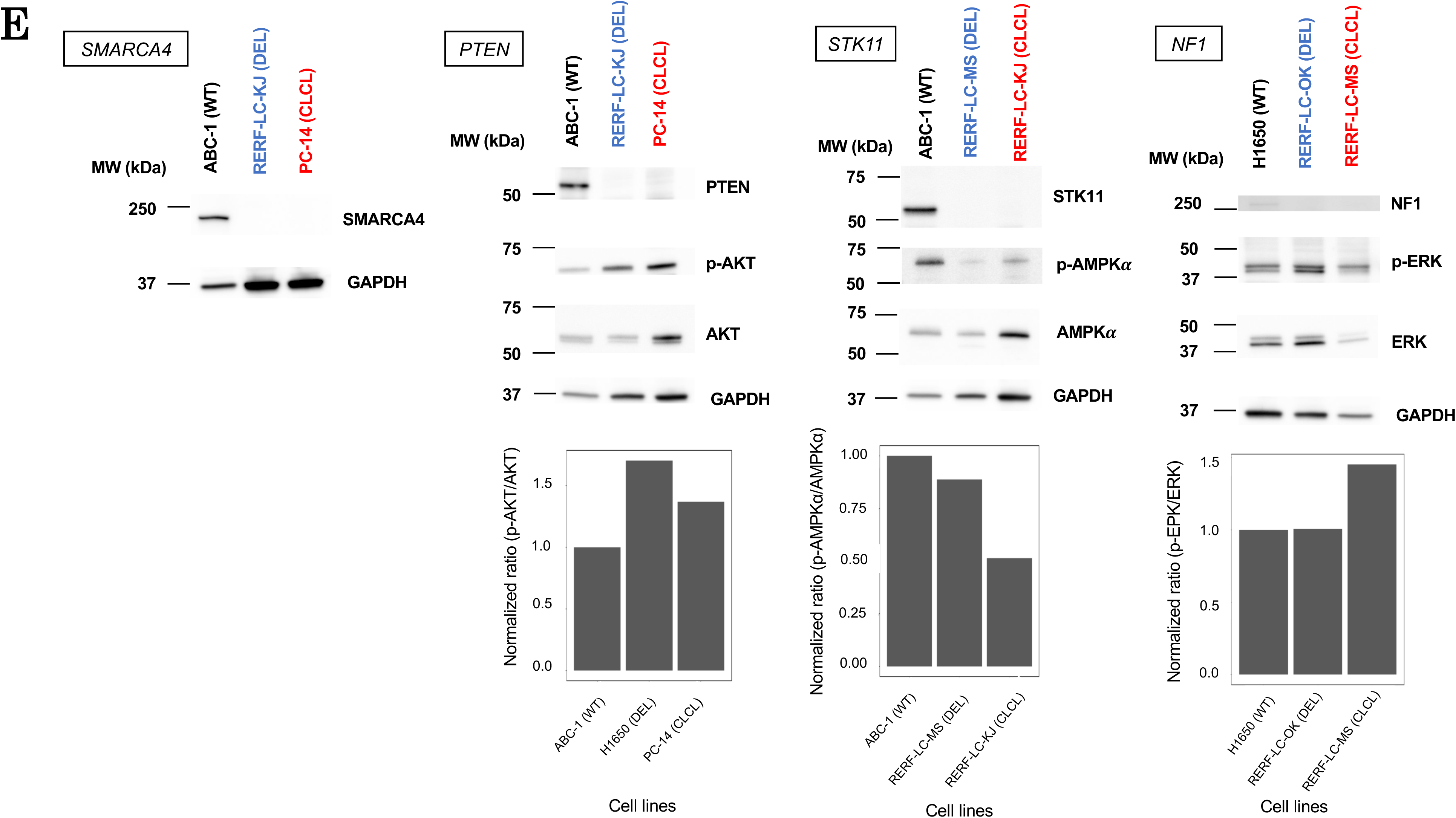
Aberrant transcriptional events caused by CLCL. (**A**) Structures of *STK11* transcripts in RERF-LC-KJ. Sequencing tags of whole-genome sequencing (PromethION) and full-length RNA-seq (MinION) were visualized by IGV. PC-14 RNA-seq was also shown as a wild-type control. (**B**) Multi-layered statuses in the *STK11* region. Patterns of whole-genome sequencing, ChIP-seq and RNA-seq tags of short read data were visualized by IGV. The status of RERF-LC-KJ and PC-9 (control) is shown. (**C**) Structures of *PTEN* transcripts in PC-14. Sequencing tags of whole-genome sequencing (PromethION) and full-length RNA-seq (MinION) were visualized by IGV. The transcripts indicated that exon 6 was skipped (black arrow). (**D**) Expression levels of *STK11, NF1, SMARCA4* and *PTEN* in 26 lung cancer cell lines. Cell lines with deleterious mutations, such as large deletions, frameshift indels and nonsense SNVs, are shown in black. Cell lines with CLCLs are also shown in black with a diagonal line. (**E**) Western blotting of genes affected by CLCLs and their downstream targets. MW: molecular weight. WT: wild type. DEL: large deletion. Bar charts indicate the normalized ratio of density of phosphorylated proteins and total proteins. Each protein is downstream of proteins encoded by CLCL genes.

We examined the epigenome marks in the regions surrounding the CLCL as represented by the ChIP-seq of H3K4me3, H3K9/14ac and RNA polymerase II. We found that chromatin normally formed the active structure at the promoter regions and that transcription was initiated normally at the correct position regardless of whether the cell line harbored the CLCL or wild-type *STK11* locus (**Fig. 4B**). However, in only the RERF-LC-KJ cells harboring the CLCL, the H3K36me3 mark disappeared in the middle of intron 1, indicating that transcriptional elongation should be disrupted exactly where the CLCL started. Illumina RNA-seq data also supported that the RNAs were abnormally spliced in the middle of intron 1 and transcribed according to the CLCL structure. The expression levels of these aberrant transcripts were measured as 2.8 rpkm. No normal transcripts were detected. However, the aberrant transcripts retained a substantial expression level, although somewhat lower than that of the wild type.

We conducted a similar analysis for the other CLCLs. For the *PTEN* gene in PC-14 (**Fig. 4C**), the CLCL resided at exon 6. As a result, this exon was completely skipped from the transcripts of *PTEN*. Accordingly, the resulting transcript should be frame-shifted and thus should be likely to lead to functional loss of the *PTEN* gene. We also examined the RNA expression levels in the *STK11, NF1, SMARCA4* and *PTEN* genes harboring CLCLs based on the Illumina RNA-seq data. The results indicated that CLCLs are generally likely to result in reduced gene expression levels (**Fig. 4D**). Nevertheless, in some cases, gene expression levels remained significant, such as the *NF1* transcripts in RERF-LC-MS cells and the *PTEN* transcripts in PC-14 cells.

To address the biological significance of the CLCLs, we examined how the CLCL-affected locus invokes changes in protein expression levels and their related signaling pathways. We conducted Western blotting analysis. As expected, we found that the proteins of STK11, NF1, SMARCA4, and PTEN were completely lost in cells harboring CLCLs in these genes (**Fig. 4E**). We further examined the activation status of the downstream proteins. The expected disruptions of the pathways were observed for all of the examined cases. PTEN suppresses the phosphorylation of AKT, and phosphorylated AKT (phospho-AKT) consequentially activates the mTOR signaling pathway^33^. Aberrant upregulation of phospho-AKT was observed, reflecting the functional loss of PTEN in PC-14 cells (PTEN-CLCL). AMPK is a gene that plays an important role in maintaining cellular homeostasis, and the phosphorylation of the AMPK protein at its alpha subunit is activated by STK11^34^. Its activation is impaired in RERF-LC-KJ cells (STK11-CLCL). The NF1 gene, which is a negative regulator of RAS^35^. Phospho-ERK, which is downstream of the RAS signaling pathway^36^, was aberrantly upregulated in RERF-LC-MS cells (NF1-CLCL). Interestingly, despite the clear protein losses of the corresponding genes in all of the cases, either by conventional aberrations or CLCLs, their consequences somewhat varied depending on the cases. For example, even though the STK11 protein similarly disappeared in both RERF-LC-MS cells (STK11-loss) and RERF-LC-KJ cells (STK11-CLCL), the enhanced ratio of phospho-AMPKα was higher in the RERF-LC-KJ cells. The effects of NF1 in RERF-LC-OK (NF1-loss) were almost undetectable, while the effects were significant in RERF-LC-MS cells (NF1-CLCL). It is possible that other pathways can sometimes complement the loss of the key protein.

### Identification of CLCLs in clinical lung cancer specimens

To examine whether CLCLs are also present in clinical cancer lung adenocarcinoma cases, we conducted similar PromethION whole-genome sequencing for the surgical specimens of nine Japanese lung adenocarcinoma patients (**Table 3** and **Supplementary Table S4**). The detected driver mutations for each patient are shown in **Table 3**. For these cases, we generated 43,953,136,203 bp sequences on average for each case (more than 10× depth; **Supplementary Table S5**). For case S10, we also sequenced normal counterparts to eliminate possible normal variations and dubious CLCLs derived from the mapping errors.

**Table 3.**
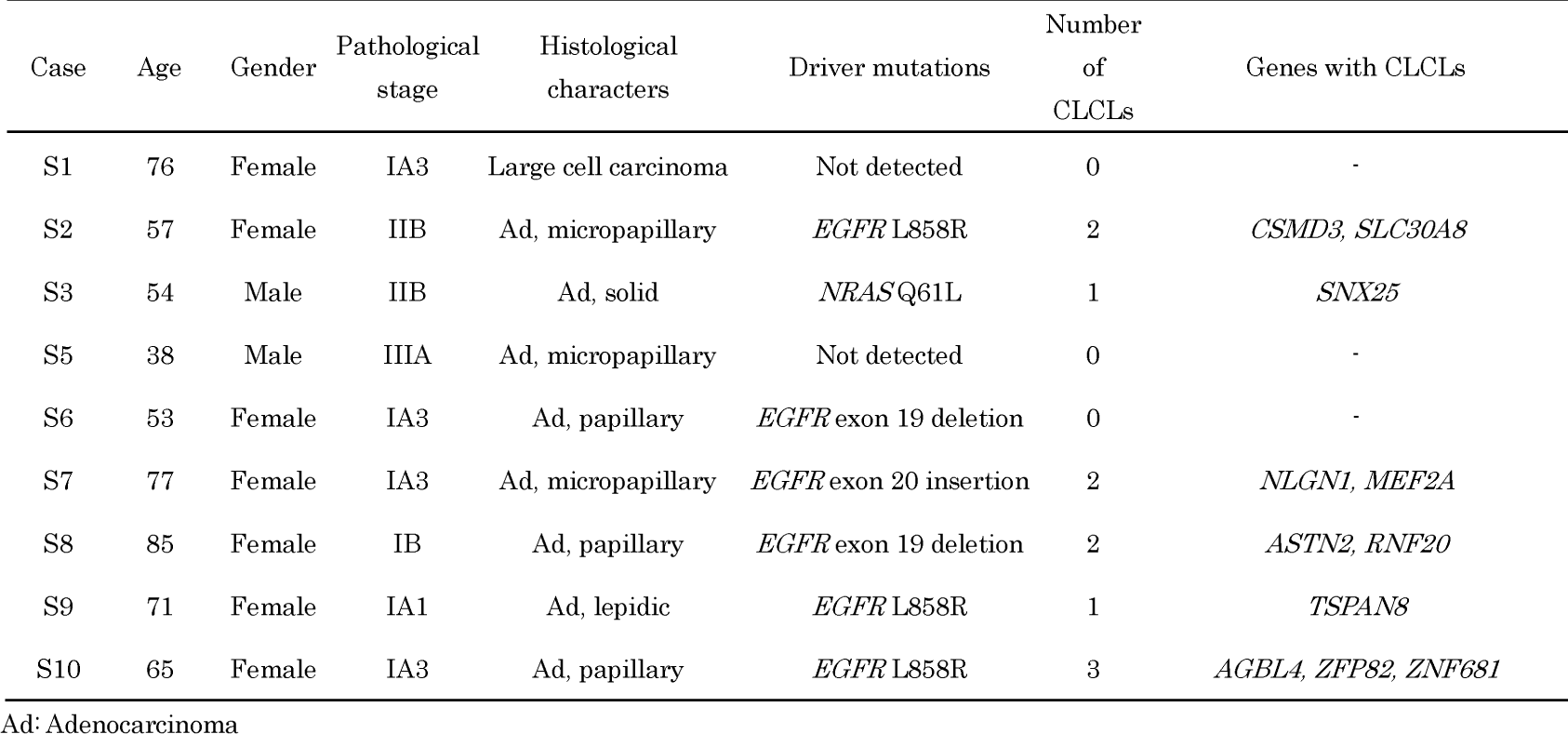
Clinical information and CLCL candidates of clinical samples.

Here, again, we successfully detected CLCLs. To our surprise, six of the nine specimens harbored at least one CLCL in their tumor genomes. Again, several key cancer genes were included. For example, we identified an *RNF20* CLCL in case S8. This patient is a female patient and had been shown to have an *EGFR* exon 19 deletion as a driver mutation. However, the other cancerous mutations remained elusive. In this case, the CLCL of the *RNF20* gene occurred as a tandem duplication between intron 2 (chr9: 101,536,324) and intron 6 (chr9: 101,544,752, **Supplementary Fig. S5**), which is very likely to lead to the functional loss of this gene. The *RNF20* gene encodes an E3 ubiquitin ligase with a tumor suppressor function, and it is frequently mutated, particularly in lung cancer^37^. In the other cases, the indications obtained for the molecular etiology underlying the carcinogenesis of the patients are summarized in **Table 3**. Further scaling the long read sequencing would be needed to more precisely identify the frequencies of the CLCLs and the preference of the genes harboring CLCLs.

### Re-evaluation of possible CLCLs based on public short read sequencing data

As the first step for scaling CLCL analysis, we attempted to utilize pre-existing Illumina short read data. We hoped that we might be able to identify CLCL candidates even from the short read sequence data. If this is possible, there are tens of thousands of whole-genome/exome sequencing datasets publicly available for the various cancer types. We were also interested in how these CLCLs had been represented by the previous short read sequences.

To identify putative CLCLs starting from the short read sequences, we employed a soft-clipping program, GenomonSV (https://github.com/Genomon-Project/GenomonSV)^38^. We selected the “split” reads as the “soft-clipped” reads and the paired-end reads, which may span the junctions of the SVs (see **Methods** for details). As the model dataset, we first analyzed the whole-genome short read data obtained for the five lung cancer cell lines that were used for the above PromethION sequencing. We could extract an average of 182 “soft-clipped” junction points in genic regions for each cell line (**Supplementary Fig. S6**). We defined tandem duplication structures as putative CLCLs in the short read data. An average of 26 genes were affected with putative CLCLs in the cell lines. We compared CLCLs detected from short reads with those from long reads (**Supplementary Table S6**). Among the CLCLs detected by PromethION, 72 % of the genes were also detected from the short read sequence data (**Fig. 5A**). However, the precision rates were limited to 25 % because of the general high rates of false-positive detection.

**Figure 5.**
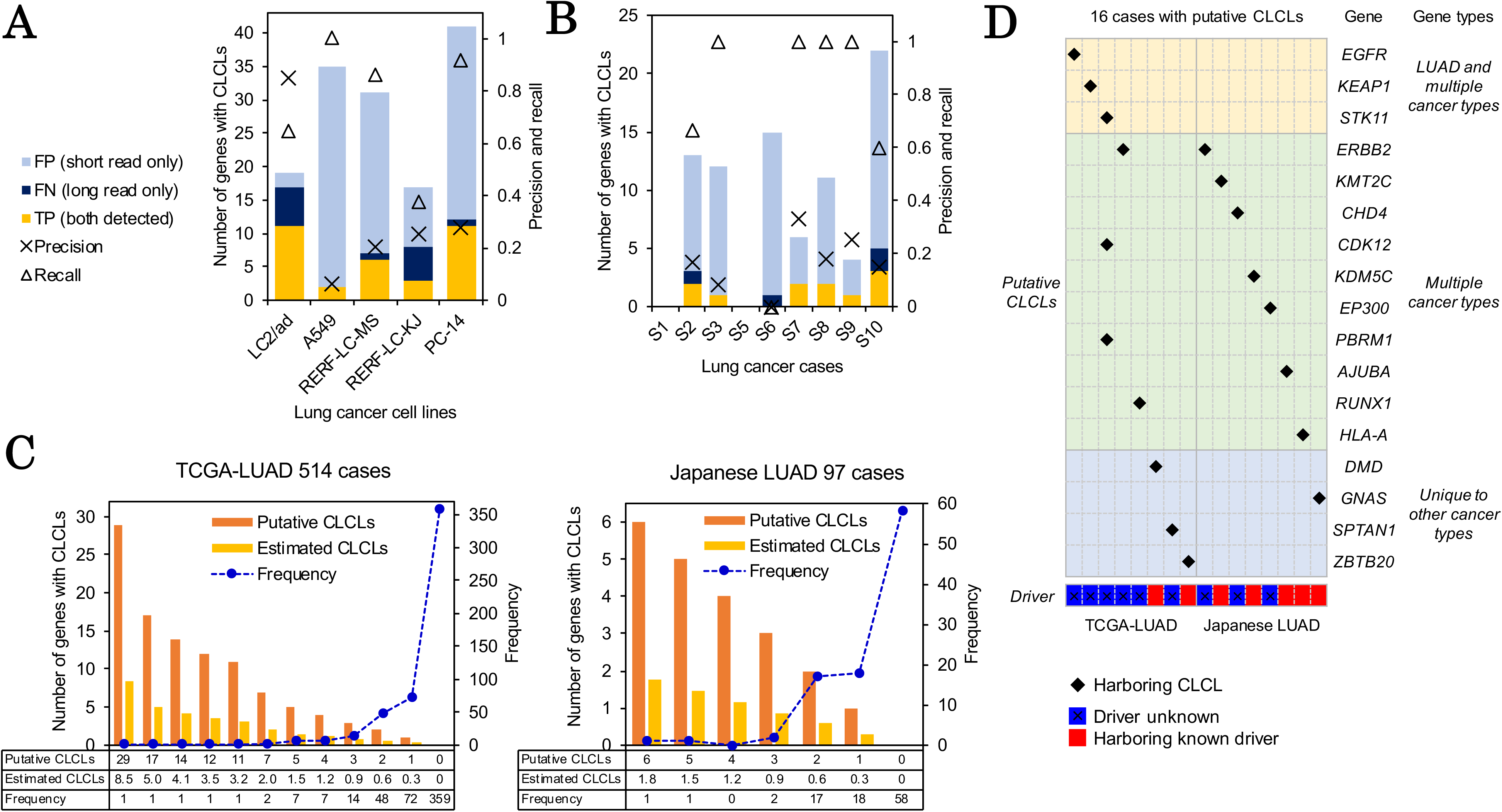
CLCL as an overlooked cancerous aberration in cancers. (**A B**) Detection of CLCLs in cell lines (**A**) and clinical samples (**B**) in short read data. The number of genes with CLCLs detected by short read data are shown in each category (TP: true positives, FN: false positives, FP: false positives; y-axis on the left side; also see **Supplementary Table S6**). The precision and recall rates are shown (y-axis on the right side). The legend and color key are shown in the margin in **A**. (**C**) The number of putative and expected CLCLs in whole-exome sequencing data. TCGA-LUAD and 97 Japanese lung adenocarcinoma datasets are shown in the left and right panels. (**D**) Putative CLCLs of cancer-related genes in 16 LUAD cases (shown as diamond). The driver mutation status of each case is shown at the bottom (blue and red). Gene types indicate driver genes that are significantly functionally altered in the indicated cancer types.

We then collected and analyzed the whole-genome short read data at a sequencing depth of approximately 63× for nine clinical cases, as shown in **Table 3**. The obtained results were inspected regarding the possible occurrence of CLCLs. We identified an average of nine genes that may be affected by putative CLCLs. As shown in **Figure 5B**, CLCLs were detected starting from the short read data at an estimated sensitivity of 73 %. However, the precision rate was supposed to be limited to 14 % for a variety of reasons inherent to the shorter read sequencing.

Despite the limited estimated precision and recall rates of CLCL detection using short read data at 21 % and 72 %, respectively, for all the cell lines and clinical samples taken together, we applied the constructed analytical pipeline to the whole-exome sequencing data of 514 TCGA lung adenocarcinoma (TCGA-LUAD)^3^ and 97 Japanese lung adenocarcinoma (Japanese LUAD)^5^ samples. We expected that the detection rate would be inherently lower, reflecting the fact that they are exome sequencing datasets. We detected a total of 269 and 50 junction points with tandem duplication structures, which are likely to correspond to the CLCLs, by soft-clipped reads from TCGA-LUAD and Japanese LUAD cases, respectively (ranging from 1 to 29 genes per case). In total, we extracted CLCL candidates from 155 (30 %) TCGA-LUAD and 39 (40 %) Japanese LUAD cases (**Fig. 5C**).

In particular, we considered whether there were any suspected cases harboring the CLCL candidates for the 299 genes that are considered to be the most relevant “cancer-associated genes”^39^. We detected 16 cases (2.6 % in 514 + 97 cases), harboring potential CLCLs in 17 genes (**Fig. 5D**). Interestingly, nine of these cases harbored no known driver mutations. For example, in the case of TCGA-49-4512 (female, nonsmoker), we identified a potential CLCL in the kinase domain of the *EGFR* gene. This duplication was previously reported^40^ and might cause aberrant activation of EGFR, thus serving as a driver mutation of this case. Importantly, this patient’s therapeutic target should be addressed by EGFR inhibitors such as afatinib^40^. Putative CLCLs associated with *ERBB2* were also detected in two other cases (both male and smoked). Aberrant duplications seemed to occur between the *ERBB2* genic region and the downstream intergenic or genic regions. Other patients were found to harbor putative CLCLs in other important tumor suppressor genes, such as *STK11* and *PBRM1*, for which mutation statuses could be utilized as putative markers for immune checkpoint inhibitors^41–43^. For these cases, the precise structure as well as the functional relevance of the putative CLCLs are still unknown, and thus, they should be subjected to detailed long read sequencing analyses.

## DISCUSSION

In this paper, we have described the identification and characterization of structural aberrations in lung cancer genomes using PromethION.

We were able to identify the precise junctions of chromosomal rearrangements and large-scale deletions relatively easily. For example, the junction points of the *CDKN2A* gene were precisely detected (**Fig. 2C**). In most cases, the proximal genes were simultaneously deleted. In LC2/ad, the deletion spanned from the *MIR31HG* and *MTAP* gene loci to the *DMRTA1* gene locus (supported by 9 reads). In A549 cells, the deletion started from the *MTAP* locus and reached the *CDKN2B* gene locus (supported by 15 reads). In PC-14, 22 genes were deleted in addition to the *CDKN2A* gene (supported by 8 reads). Several studies have reported that *CDKN2A* codeleted genes are involved in the hidden molecular features of cancers. The *MTAP* gene encodes 5-methylthioadenosine phosphorylase, which is associated with the purine and methionine salvage pathways, located to the adjacent region of the *CDKN2A* gene and frequently codeleted in cancers. MTAP-deficient cancers are known to acquire vulnerability to arginine methyltransferase PRMT5 depletion, which may be a novel target of an anticancer drug^44–46^. It is important to determine the precise junction by the long read approach to completely understand what genes or regions are affected by these genomic aberrations.

We also identified the 8 Mb amplification for the *MYC* gene locus in LC2/ad cells. We attempted to further characterize it and found that even the latest long read sequencing technologies could not comprehend the precise mutation pattern in this locus. Eight Mb may have been too large, and the internal rearrangement may have been too complicated, although this is the only region where we could not reassemble the structure. Interestingly, we found no sequences suggesting aberrations occurring within the internal region of the *MYC* gene locus itself. Outside of the *MYC* gene, at least four break points were detected by nanopore reads, which were further confirmed by the Illumina short reads (**Fig. 2D**). There may be a unique selective pressure exerted on this gene specifically, retaining the gene function itself intact, at the same time, enhancing its gene expression.

Most importantly, in this study, we unexpectedly identified a unique aberration pattern, the CLCL. We found that CLCLs exist even in pivotal cancer-related genes, such as the *STK11, NF1, SMARCA4,* and *PTEN* genes. Recent papers have reported that immune checkpoint inhibitors are less effective for lung cancers with *STK11* mutations^41,42^. Therefore, the therapeutic strategy for each patient would be different depending on whether there is a mutation in the *STK11* gene or not. Additionally, CLCLs were identified in the *PTPN13, RPTOR*, and *RHEB* genes in LC2/ad cells (**Table 2**). *PTPN13* encodes a protein tyrosine phosphatase. The *RPTOR* and *RHEB* genes are members of the mTOR signaling pathway. The functional loss of these genes should be related to tumorigenesis and malignancy of the cancer, although further studies will be needed to clarify the relationship between those aberrations and the molecular etiology of the cancers in more detail.

We could also analyze the causes and consequences of genomic SVs. In total, 67 % of the CLCLs had at least one junction in LINE, SINE and LTR regions, suggesting that transposable elements were likely to contribute to the formation of CLCLs. Using epigenome and transcriptome data, we also showed that CLCLs led to the formation of abnormal transcripts and functional loss of their encoded proteins in most cases.

We further conducted long read sequencing of clinical samples. We successfully demonstrated that CLCLs occur in the *in vivo* genomes of lung adenocarcinoma patients. In six cases, at least one CLCL was detected in the genes of important functions, giving the complementary therapeutic indication for the patients. Finally, we reanalyzed short read sequencing data from clinical samples that were previously published^3,5^, particularly focusing on detecting CLCLs in cancer-related genes, such as the driver genes of lung adenocarcinoma. Although the current precision rate is limited, the recall rate was reasonably high. We believe it is important to subject those cases to further detailed long read sequencing. We suggest that CLCLs occurring in cancer genomes might have important roles in the phenotypic features of cancers, including responses to anticancer drugs.

Lastly but not less importantly, we found that the visualization of the detected mutation is also important. So-called “genome graph” databases should play an indispensable role in representing the diverse nature of cancer genomes and thus further enhance the accuracy of future genome analyses^47,48^. This is the first study that has utilized PromethION sequencing for cancer genomics. Obviously, further improvements in the sequencing method itself, coupled with refinements of the computational tools, are needed to reach further goals. Indeed, this study may have presented more questions than answers. In that sense, this is only the first study paving the way towards a more comprehensive understanding of the complicated genomic aberrations of cancers and further in-depth study of their biology.

## MATERIAL AND METHODS

### Cell lines and clinical samples

The Lung adenocarcinoma cell lines LC2/ad, A549, RERF-LC-KJ, RERF-LC-MS, and PC-14 were cultured as previously described^23^. Cell pellets were washed with cold PBS and cryopreserved.

Clinical samples were obtained with the appropriated informed consent at the National Cancer Center Japan. Surgical specimens from 10 patients were pathologically checked, and one was removed because of low tumor content (**Supplementary Fig. S7** and **Supplementary Table S4**). All nine patients were diagnosed with primary lung cancer, including eight adenocarcinomas and one large cell carcinoma (**Table 3**). Fresh frozen surgical specimens were used to extract genomic DNA (gDNA) and total RNA as described below.

### Whole-genome sequencing using MinION

High-molecular-weight (HMW) gDNA was extracted from the lung cancer cell lines LC2/ad and A549 with Smart DNA prep(a) kit (Analytikjena). In the case of LC2/ad, WGS data were produced from 1D sequencing (SQK-LSK108), 1D^2^ sequencing (SQK-LSK308), rapid sequencing (RAD003), and in the case of A549, WGS data were produced from only 1D^2^ sequencing. In summary, 4 µg HMW gDNA was used for 1D sequencing and DNA repair, end-prep, and adapter ligation were conducted. DNA repair was performed using NEBNext FFPE DNA Repair Mix (M6630, NEB). End-prep was performed using NEBNext Ultra II End Repair/dA-Tailing Module (E7546L, NEB). Adapter ligation was performed using NEBNext Blunt/TA Ligase Master Mix (M0367L, NEB) and Ligation Sequencing Kit 1D (SQK-LSK108, Oxford Nanopore Technologies). In summary, 1D^2^ sequencing, 4 or 5 µg HMW gDNA was used as input, and DNA repair, end-prep, first adapter ligation, and second adapter ligation were conducted. DNA repair and end-prep were the same protocol as the 1D sequencing. First and second adapter ligations were performed using NEBNext Blunt/TA Ligase Master Mix and Ligation Sequencing Kit 1D^2^ (SQK-LSK308, Oxford Nanopore Technologies). DNA purifications in each step of 1D and 1D^2^ sequencing were performed using Agencourt AMPure XP (A63882, Beckman Coulter). In summary, 15 µl gDNA was used for rapid sequencing, and a Rapid Sequencing Kit (RAD003) was used.

### Whole-genome sequencing using PromethION

The HMW gDNA extraction method was the same as MinION sequencing for LC2/ad, A549, and RERF-LC-MS. Forty-eight microliters of 1.5 or 2 µg gDNA plus nuclease free water (NFW), 3.5 µl of NEBNext FFPE DNA Repair Buffer, 2 µl of NEBNext FFPE DNA Repair Mix (M6630, NEB), 3.5 µl of NEBNext Ultra II End Prep Reaction Buffer, and 3 µl of NEBNext Ultra II End Prep Enzyme Mix (E7545L, NEB) were mixed gently in 1.5 ml Eppendorf tube. After spinning down, the sample was incubated at 20 °C for 5 minutes and 65 °C for 5 minutes. Then, 60 µl of AMPure XP beads (A63882, Beckman Coulter) was added to the tube and the tube was mixed by flicking. The sample was incubated for 5 minutes at room temperature (R.T.) using a rotator mixer. After spinning down, the tube was set on a magnetic stand, and the supernatant was pipetted off. The beads were washed twice with 200 µl of 70 % ethanol. The tube was spun down and back on the magnet. Any residual ethanol was pipetted off, and the tube was dried for 30 seconds. The tube was removed from the magnetic stand, the pellet was resuspended in 61 µl NFW, and the tube was incubated for 2 minutes. The tube was set on a magnet, 60 µl of the sample was used in the next step, and 1 µl was used for quality check by Qubit. 60 µl of end-prepped DNA, 25 µl of Ligation Buffer (LNB), 10 µl of NEBNext Quick T4 DNA Ligase (NEB, E6056S), and 5 µl of Adapter Mix (AMX) were mixed in a 1.5 ml Eppendorf tube. After spinning down, the sample was incubated at R.T. for 10 minutes. Then, 40 µl of AMPure XP beads was added to the tube and mixed by flicking. The sample was incubated for 5 minutes at R.T. using a rotator mixer. After spinning down, the tube was set on the magnetic stand. The supernatant was pipetted off, and the beads were washed twice with 250 µl of Fragment Buffer (LFB). The sample was spun down, and the tube was replaced on the magnet. Any residual supernatant was pipetted off, and the tube was dried for 30 seconds. The tube was removed from the magnet, and the pellet was resuspended in 25 µl of Elution Buffer (EB). The sample was incubated at R.T. for 10 minutes. Next, 24 µl of the sample was used in the next step, and 1 µl was used for quality check by Qubit. A total of 46 µl of Flush Tether (FLT) was added directly to the tube of PromethION Flush Buffer (PFB), and the solution was mixed by pipetting (Priming Mix). Then, 800 µl of Priming Mix was loaded onto the PromethION flow cell. Next, 75 µl of SQB, 51 µl of LB, and 24 µl of the DNA library were mixed in a 1.5 ml Eppendorf tube in 5 minutes (Loading Library). Then, 200 µl of Priming Mix was loaded onto the flow cell, and 150 µl of Loading Library was loaded onto the flow cell. We started the PromethION run. The LNB, AMX, LFB, EB, FLT, PFB, SQB, and LB are in Ligation Sequencing Kit 1D (SQK-LSK109, Oxford Nanopore Technologies). From RERF-LC-KJ, PC-14, and lung adenocarcinoma clinical samples, HMW gDNA was extracted with the MagAttract HMW DNA Kit (Qiagen). For the LC2/ad cells, we performed five runs, and each throughput was 10.4 Gb, 9.2 Gb, 33.0 Gb, 28.5 Gb, and 19.3 Gb, respectively.

### Full-length transcriptome sequencing using MinION

Full-length transcriptome analysis using MinION was performed as previously described^49^. RNA was extracted from lung cancer cell lines using the RNeasy Mini kit (Qiagen). The extracted RNA was converted to cDNA using SMART-seq v4 Ultra Low input RNA kit (Takara). Then, we used cDNA as input for 1D^2^ MinION sequencing.

### Computational analysis of long read sequencing data

MinION fast5 data were basecalled using albacore 2.0.2 and converted fastq files. PromethION fast5 data were basecalled using guppy and converted fastq files. Our MinION and PromethION data set were mapped to the human reference genome, hg38, using Minimap2 (with the “-ax map-ont” option, 2.9-r720 version). MinION 1D^2^ sequencing outputs two types of fastq files, 1D and 1D^2^. 1D means that reads were generated using single strand information. 1D^2^ reads integrate the double strand information. There were some overlapping reads between 1D files and 1D^2^ files. Therefore, reads used as 1D^2^ were removed from 1D files. In addition, there were some overlapping reads in the 1D^2^ files. These reads were removed from the 1D^2^ files and used as 1D reads.

### Detection of driver mutations for cell lines in long read data

We detected known driver mutations by IGV. For the point mutations, we directly explored the known positions of the mutations. For the driver mutation of LC2/ad cells, the *CCDC6-RET* fusion gene, we explored the reads split-aligned to both *RET* and *CCDC6* genes in the reference genome and extracted the alignments with SAM format from IGV. Then, we extracted the information of split alignment (chromosome, position of reference, read strand, and position of read) from the file and sorted the information by the position of reads. We filtered out reads with small MAPQ (< 30) and counted the number of supporting reads.

### Detection of structural variants from long read data

To detect gene rearrangements and CLCLs, we used the information of split alignments. First, we mapped sequencing data (fastq) to the human reference genome, hg38. Then, we extracted reads with split alignment from mapping data (bam) using the command “samtools view -f 2048”. We filtered out reads with multiple hits (flag: 256). Then, we extracted the information of split alignments (chromosome, position of reference, read strand, and position of read) from the file and sorted the information by the position of reads. We filtered out reads with low MAPQ (< 30) from the dataset. We extracted junction candidates of gene rearrangements and CLCLs (tandem duplications and inversions) considering the position of reads (removing junctions with large differences in read position, >300 bp), annotated the junctions by genes from DBKERO (http://kero.hgc.jp/) and merged the junctions less than 50 bp from the junctions of CLCL candidates. The threshold for the number of reads supporting the junctions was four in the clinical samples and five in the cell lines. We removed junctions with less than 2,000 bp between the junctions. Finally, we checked the structure of the candidates for gene rearrangements and CLCLs by IGV and manually removed questionable rearrangements and CLCLs. To detect deletions, we used the information of split alignments and CIGAR strings in SAM format files. We extracted the information of split alignments in the same way as the detection of gene rearrangements and CLCLs. We extracted the CIGAR strings from reads with primary alignments using SAMtools (flags: 0 or 16). Then, we detected deletions over 2,000 bp.

### Optical mapping using the Saphyr system

Optical mapping analysis using the Saphyr system (Bionano Genomics) was performed for LC2/ad cells. Briefly, HMW DNAs were isolated from frozen cells using a Bionano Prep kit (Bionano Genomics) and measured by Qubit BR assay (Invitrogen). The extracted DNAs were fluorescently labeled with DLE-1 using a Bionano DLS kit (Bionano Genomics). Data were collected on the Saphyr instrument (Bionano Genomics). The figures were created by Bionano Access (version 1.3.0, Bionano Genomics).

### Western blotting

We performed Western blotting as described previously to quantify proteins from genes with CLCL structures^50^. For Western Blotting, we performed protein extraction and quantification. We used a Pierce BCA Protein Assay kit (23225, Thermo Fisher SCIENTIFIC) and prepared proteins of 1 mg/ml. In summary, we performed electrophoresis of proteins, membrane transferring, blocking, reaction of the first antibody, reaction of the second antibody, and visualization of bands. We conducted electrophoresis using a tank and used 10 or 15 µl of proteins as input. For blocking, we used 4 % BSA blocking buffer. In the reaction of the first antibody, we used LKB1 (27D10) Rabbit mAb (3050S, CST) for STK11, phospho-AMPK-alpha(Thr172) Antibody (2535S, CST) for phospho-AMPKα, AMPK-alpha Antibody Rabbit mAb (2603S, CST) for AMPKα, Anti-Neurofibromin (NF1) (rabbit polyclonal IgG) (07-730, upstate) for NF1, Phospho-p44/42 MAPK (Erk1/2) (Thr202/Thr204) (E10) Mouse mAb (9106S, CST) for phospho-ERK, p44/42 MAPK (Erk1/2) Antibody (9102S, CST) for ERK, Brg1 (D1Q7F) Rabbit mAb (49360, CST) for SMARCA4, PTEN (138G6) Rabbit mAb (CST, 9559S) for *PTEN* protein, Phospho-Akt (Ser473) (D9E) XP ® Rabbit mAb (4060S, CST) for phospho-AKT, and Akt (pan) (11E7) Rabbit mAb (4685S, CST) for AKT. As a control, we used GAPDH and GAPDH (14C10) Rabbit mAb (2118S, CST) as an anti-body. In the reaction of the second antibody, we used an antibody corresponding to the animal that generated the first antibody. We used Anti-rabbit IgG, HRP-linked antibody (7074S, CST) for rabbit and Anti-mouse IgG, HRP-linked antibody (7076S, CST) for mouse. For the visualization of bands, we used ImageQuant LAS 4000 mini (GE Healthcare). The normalized ratio indicated the fraction of relative density of phosphorylated protein to control protein and relative density of total protein to control protein, and we set the normalized ratio of the wild type as one (**Fig. 4E**). The density of the bands was calculated by ImageJ software^51^.

### Whole-genome short read sequencing of clinical samples

Genomic DNA was extracted from surgical specimens using the MagAttract HMW DNA Kit (QIAGEN). Whole-genome sequencing libraries were constructed using the TruSeq Nano DNA Library Prep kit (Illumina) and sequenced by NovaSeq according to the manufacturers’ instructions. In summary, 100 ng gDNA was used for library preparation as input, and DNA fragmentation, end repair, adenylation of the 3’ end, adapter ligation, and condensation of DNA fragments were conducted. We performed DNA fragmentation using Covaris (Covaris) and used a protocol for 350 bp of insert size. After preparing the library, we denatured the library with NaOH and then started the NovaSeq run.

### Analysis of short read sequencing data

Whole-genome short read sequences were mapped to the human reference genome (hg38) using BWA-MEM (version 0.7.15). After mapping, sorted BAM files were created, and PCR duplicates were marked by SAMtools. For detection of driver mutations in the ten clinical samples, point mutations and variations were called using GATK Mutect2 (version 4.0.12.0).

For transcriptome and epigenome analysis of the cell lines, we used RNA-seq and ChIP-seq data that were previously obtained (DRA001846 and DRA001860) and mapped to the reference genome hg19. IGV was used for visualization.

To detect SV junctions, GenomonSV (version 2.6.1) was used with paired-end read sequencing data as listed; 5 whole-genome datasets from cancer cell lines (DRA001859; 101PE, Illumina HiSeq2500), nine whole-genome datasets from Japanese lung cancer patients (150PE, Illumina NovaSeq), 514 whole-exome datasets from TCGA-LUAD and 97 whole-exome datasets from Japanese lung adenocarcinoma patients (JGAS00000000001; 76PE, Illumina GAIIx). After conducting Genomon (version 2.6.1) with the recommended parameters (https://genomon.readthedocs.io/ja/latest/; reference: hg19), GenomonSV filt was performed with the options “--min_junc_num 1” and “--non_matched_control_junction” with control panels that were constructed from ICGC/TCGA data. For clinical samples harboring matched normal data, we also set the option “--matched_control_bam”. For the cell lines, matched normal data of case S1 were used for normal control data. After GenomonSV filt, we eliminated SV candidates with tumor VAF < 0.05 and a distance of less than 2 kb between the junctions. We analyzed at least one SV junction within genic regions. For validation of SVs, junction points were compared between long read and short read data, allowing 100-bp margins after performing conversion of the reference genome version (https://genome.ucsc.edu/cgi-bin/hgLiftOver).

Genes with putative CLCLs are defined as those affected by SV junctions with a tandem duplication structure. To evaluate the detection power of short read data, genes with CLCL (and putative CLCLs) were classified into three categories: true positives detected in both long read and short read data; false positives detected in short read data only; and false negatives detected in long read data only. The precision and recall rates of CLCL detection of short read data were calculated using the number of true positives, false positives and false negatives. For CLCLs in cancer-related genes, we checked 299 cancer-related genes that were previously reported as cancer driver genes. The cancer driver genes were classified in gene types as follows: genes significant in lung adenocarcinoma and pan-cancer types as “LUAD and multiple cancer types”, genes significant in multiple and pan-cancer types as “Multiple cancer types”, and genes significant in other cancer types than lung adenocarcinoma as “Unique to other cancer types”^39^. For known driver mutation status, 13 genes (*EGFR, KRAS, BRAF, HRAS, NRAS, RET, MAP2K1, ALK, ROS1, ERBB2, MET, NF1* and *RIT1*) were examined in the 16 cases by cBioPortal (Lung adenocarcinoma; TCGA; PanCancer Atlas)^52–54^.

## DATA ACCESS

All sequencing data of cell lines were published in the DNA Data Bank of Japan (DDBJ) under the accession numbers, DRA007423 (DRX143541, DRX143542, DRX143543, DRX143544), DRA007941, DRA008154 and DRA008295. The data were also deposited in DBKERO (https://kero.hgc.jp)^55^. Sequencing data of clinical samples were deposited at the Japanese Genotype-phenotype Archive (JGA, http://trace.ddbj.nig.ac.jp/jga), which is hosted by the National Bioscience Database Center (NBDC) and DDBJ, under the accession numbers, JGAS00000000065 (JGAD00000000252 and JGAD00000000253).

## ACKNOWLEDGEMENTS

We thank K. Imamura, K. Abe, M. Kimura, H. Wakaguri, Y. Kuze and T. Horiuchi for their technical assistance. The Bionano data were generated at Bionano Genomics and K. Hong and A. Pang collected and analyzed the data. This work was supported by AMED P-CREATE Grant Number JP19cm0106539. This work was also supported in part by The National Cancer Center Research and Development Fund (29-A-6). This work was also supported by JSPS KAKENHI Grant Number 16H06279. The results shown in this study are in part based on data generated by the TCGA Research Network (https://www.cancer.gov/tcga). The super-computing resource was provided by Human Genome Center, the University of Tokyo (http://sc.hgc.jp/shirokane.html).

## AUTHOR CONTRIBUTIONS

Y. Sakamoto, A.S. and Y. Suzuki designed the study. Y. Sakamoto, L.X., and M.S. performed sequencing experiments. Y. Sakamoto, T.K., M.K., Y. Shiraishi and A.S. contributed computational analysis of sequencing data. Y. Sakamoto, Y.K., A.O. and S.K. conducted Western blotting. Y. Shimada, N.M. and T.K. contributed and analyzed clinical specimens. K.T., S.K., T.K. and Y. Suzuki interpreted the findings and supervised the study. Y. Sakamoto, A.S. and Y. Suzuki wrote the manuscript. All authors approved the final version of the manuscript.

## SUPPLEMENTARY FIGURES AND TABLES

**Supplementary Figure S1 General statistics of MinION and PromethION in four cell lines**

**Supplementary Figure S2 IGV images of large deletions detected by MinION and PromethION**

**Supplementary Figure S3 A novel genomic rearrangement in RERF-LC-KJ**

**Supplementary Figure S4 Pipeline for the detection of CLCLs**

**Supplementary Figure S5 An example CLCL from the clinical samples**

**Supplementary Figure S6 SV junctions detected by GenomonSV**

**Supplementary Figure S7 Representative histological images of clinical lung cancer specimens**

**Supplementary Table S1 Summary of lung cancer cell lines**

**Supplementary Table S2 Sequencing statistics of MinION and PromethION**

**Supplementary Table S3 Candidate novel fusion genes**

**Supplementary Table S4 Histopathological information on lung cancer clinical samples**

**Supplementary Table S5 General statistics of PromethION in lung cancer clinical samples**

**Supplementary Table S6 Numbers of genes affected by CLCLs in cell lines and clinical samples**

