## Supplementary Figures and Tables for "Long read sequencing reveals a novel class of structural aberrations in cancers: identification and characterization of cancerous local amplifications"

Supplementary Figures S1 - S7

Supplementary Tables S1- S6

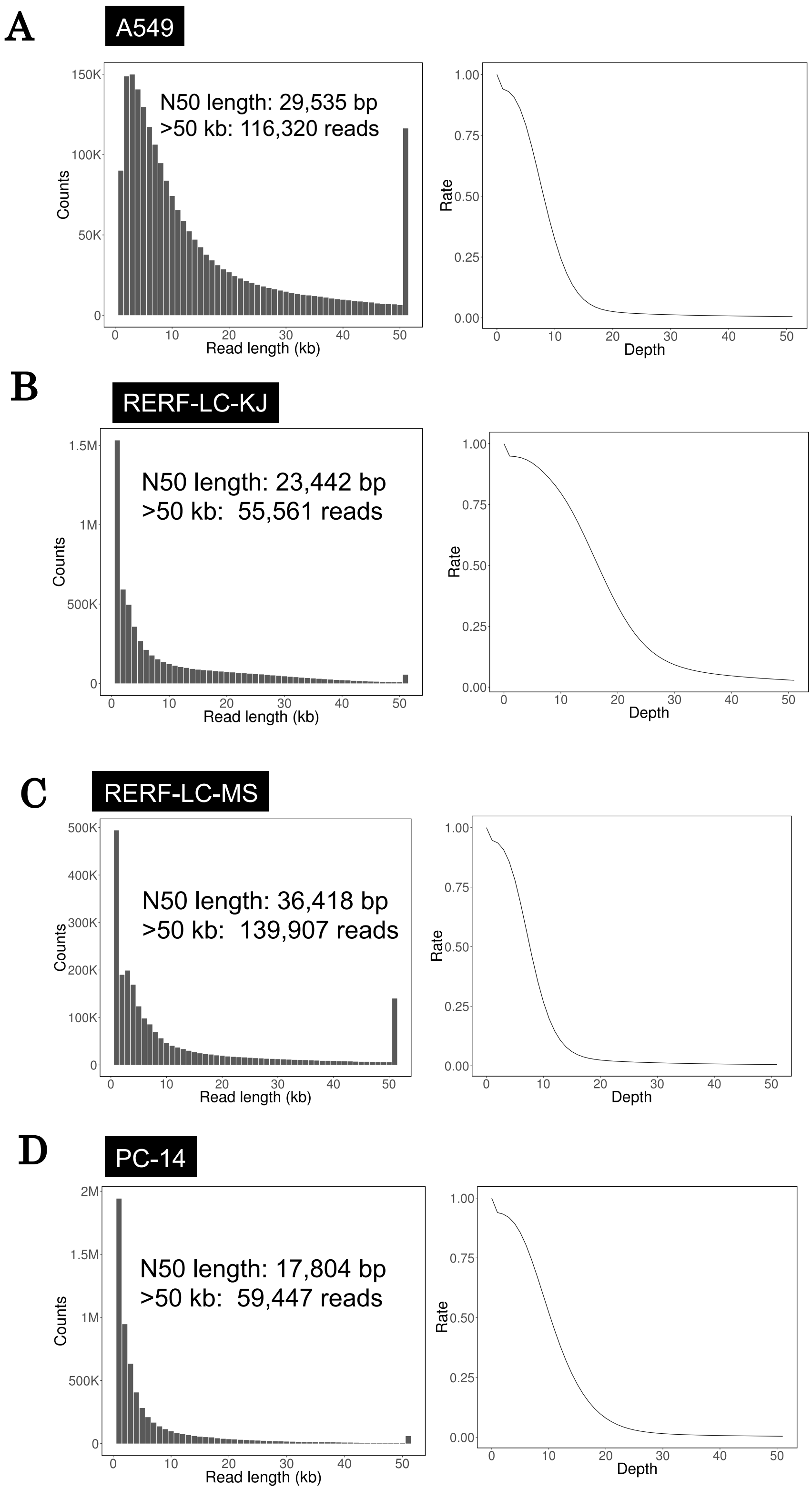
A
A549
N50 length: 29,535 bp
>50 kb: 116,320 reads
Counts
Read length (kb)
Rate
Depth
B
RERF-LC-KJ
N50 length: 23,442 bp
>50 kb: 55,561 reads
Counts
Read length (kb)
Rate
Depth
C
RERF-LC-MS
N50 length: 36,418 bp
>50 kb: 139,907 reads
Counts
Read length (kb)
Rate
Depth
D
PC-14
N50 length: 17,804 bp
>50 kb: 59,447 reads
Counts
Read length (kb)
Rate
Depth

**E**

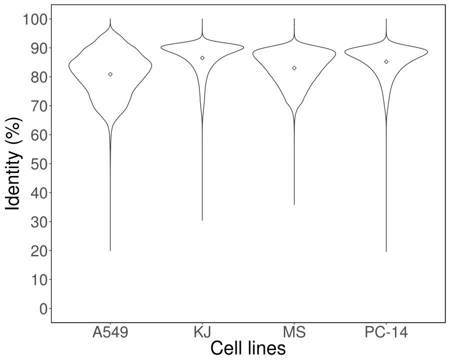


**Supplementary Figure S1 General statistics of MinION and PromethION sequencing in four cell lines.**

(**A**-**D**) Length distribution of whole-genome sequencing reads (left) and cumulative depth curve of each cell line (right). In the length distribution, the number of reads over 50 kb and N50 length of reads are displayed. (**A**) MinION sequencing of A549 cells. (**B**) PromethION sequencing of RERF-LC-KJ cells. (**C**) PromethION sequencing of RERF-LC-MS cells. (**D**) PromethION sequencing of PC-14 cells. (**E**) Percent sequence identity of the mapped reads in each cell line. KJ: RERF-LC-KJ. MS: RERF-LC-MS. The points indicate the average of sequence identity of each cell line.

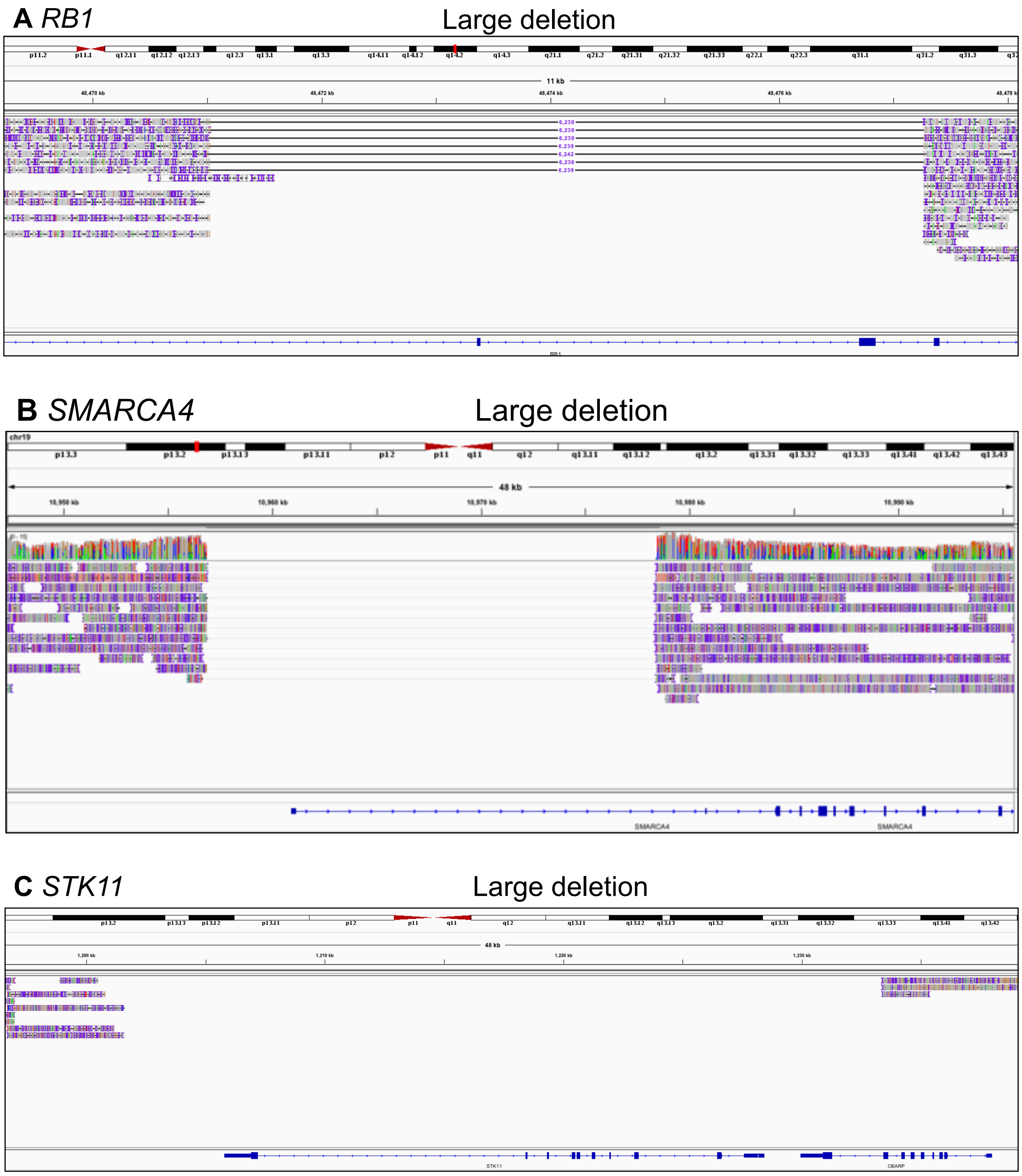


**Supplementary Figure S2 IGV images of large deletions detected by MinION and PromethION.**

**(A)** A large deletion of *RB1* in RERF-LC-KJ. **(B)** A large deletion of *SMARCA4* in RERF-LC-KJ. **(C)** A large deletion of *STK11* in RERF-LC-MS. These deletions were previously detected using Illumina sequencing.

RERF-LC-KJ

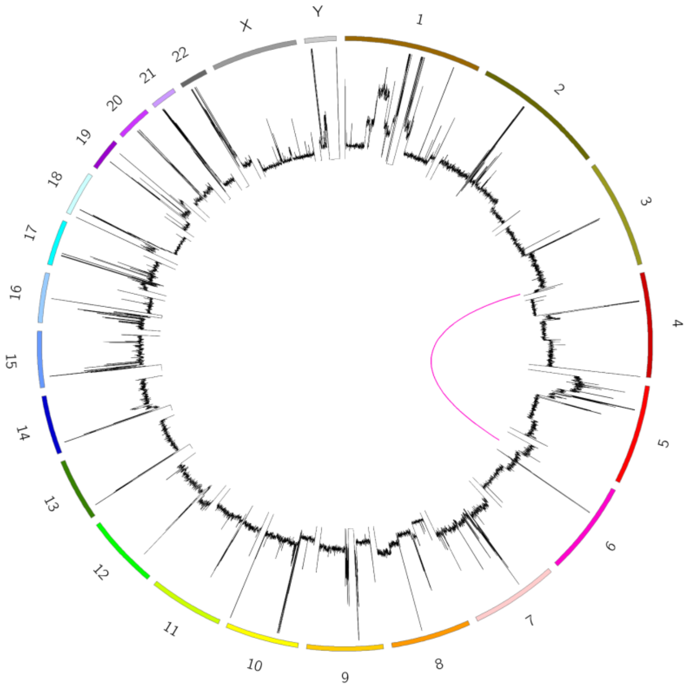


**Supplementary Figure S3 A novel genomic rearrangement in RERF-LC-KJ.**

A genomic translocation in RERF-LC-KJ is shown in the circus plot. The detailed information of this rearrangement is shown in **Supplementary Table S3**.

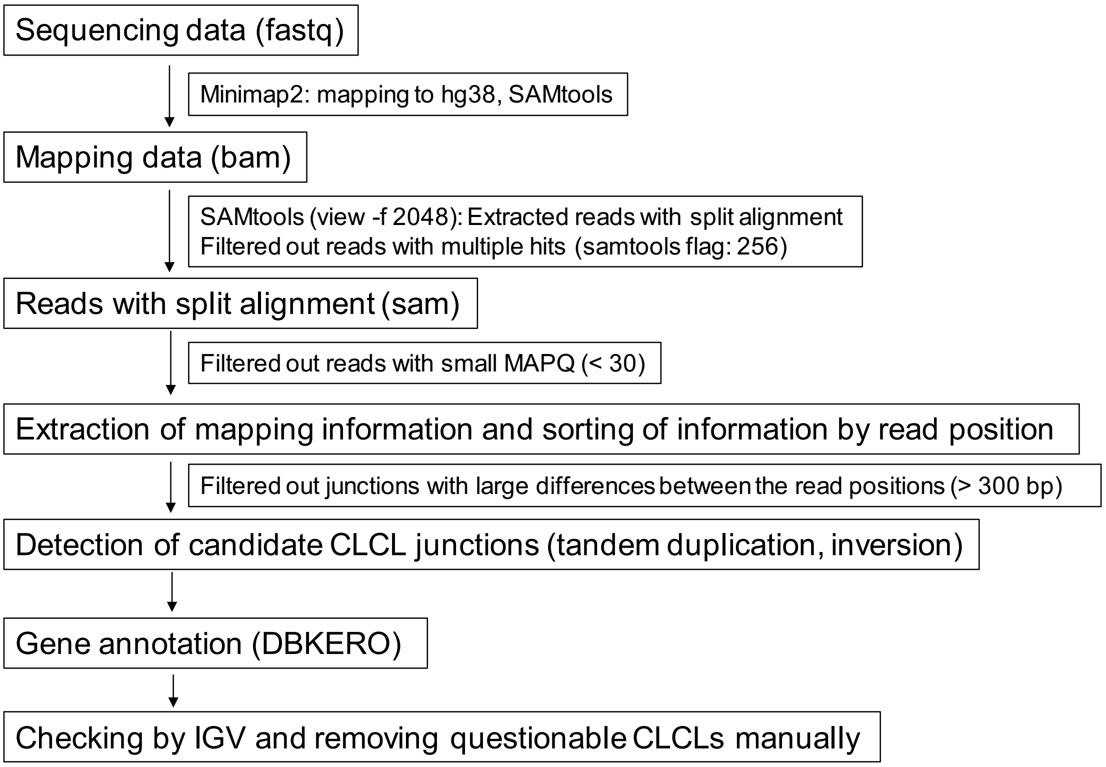


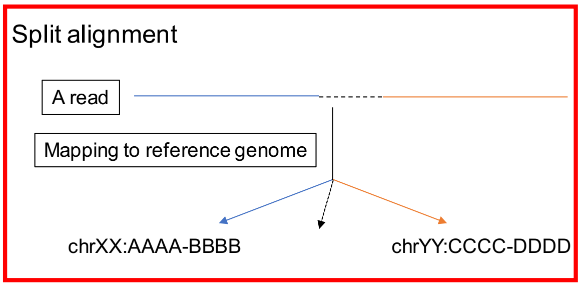


**Supplementary Figure S4 Pipeline for the detection of CLCLs.**

The detailed procedure is shown in **Material and Methods**. Briefly, long read sequences in a fastq file were mapped to the reference genome hg38 using Minimap2. Reads with split alignment were extracted using the SAMtools view with the option "-f 2048". Reads with multiple hits (flag: 256) and <30 MAPQ were filtered out. Junction candidates for SVs were extracted considering the position of the reads and merged allowing 50-bp margins. The junctions supported by ≥5 or ≥4 reads were extracted as SVs in cell lines and clinical samples, respectively. Candidates with less than 2,000 bps between the junctions were discarded. The structures of the candidates of SVs were visualized by IGV and checked manually. Split alignment means that a read can be split and the subreads can be mapped to different positions in the reference genome.

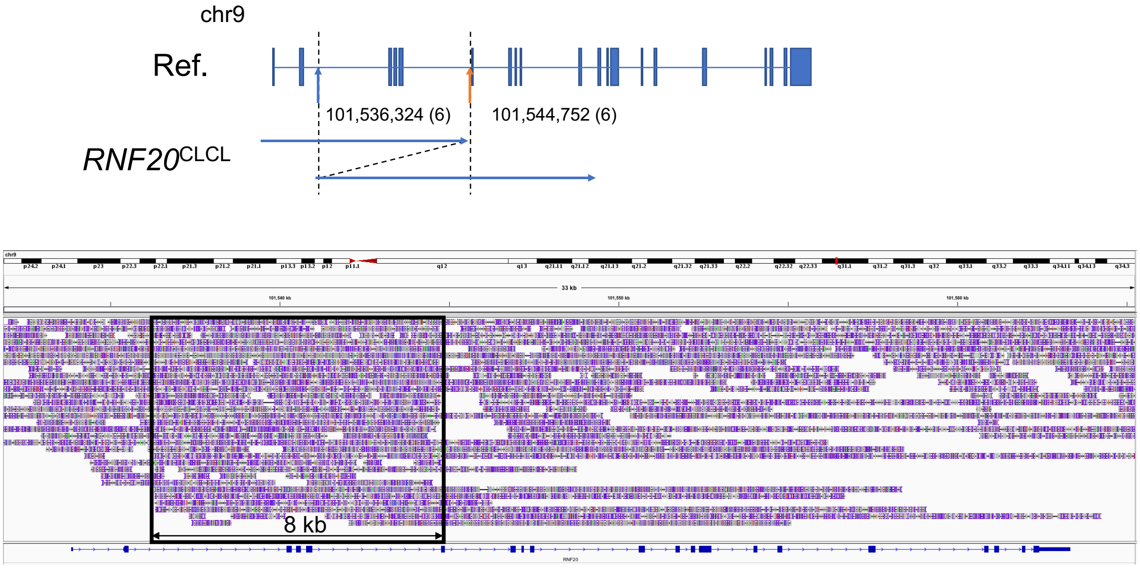


**Supplementary Figure S5 An example CLCL from the clinical samples.**

*RNF20* CLCL in case S8. An aberrant structure of *RNF20* is shown in the upper panel. The structure was a tandem duplication between the junctions (blue arrow and yellow arrow). IGV visualization is represented in the lower panel. A CLCL structure (approximately 8 kb) is indicated in the box.

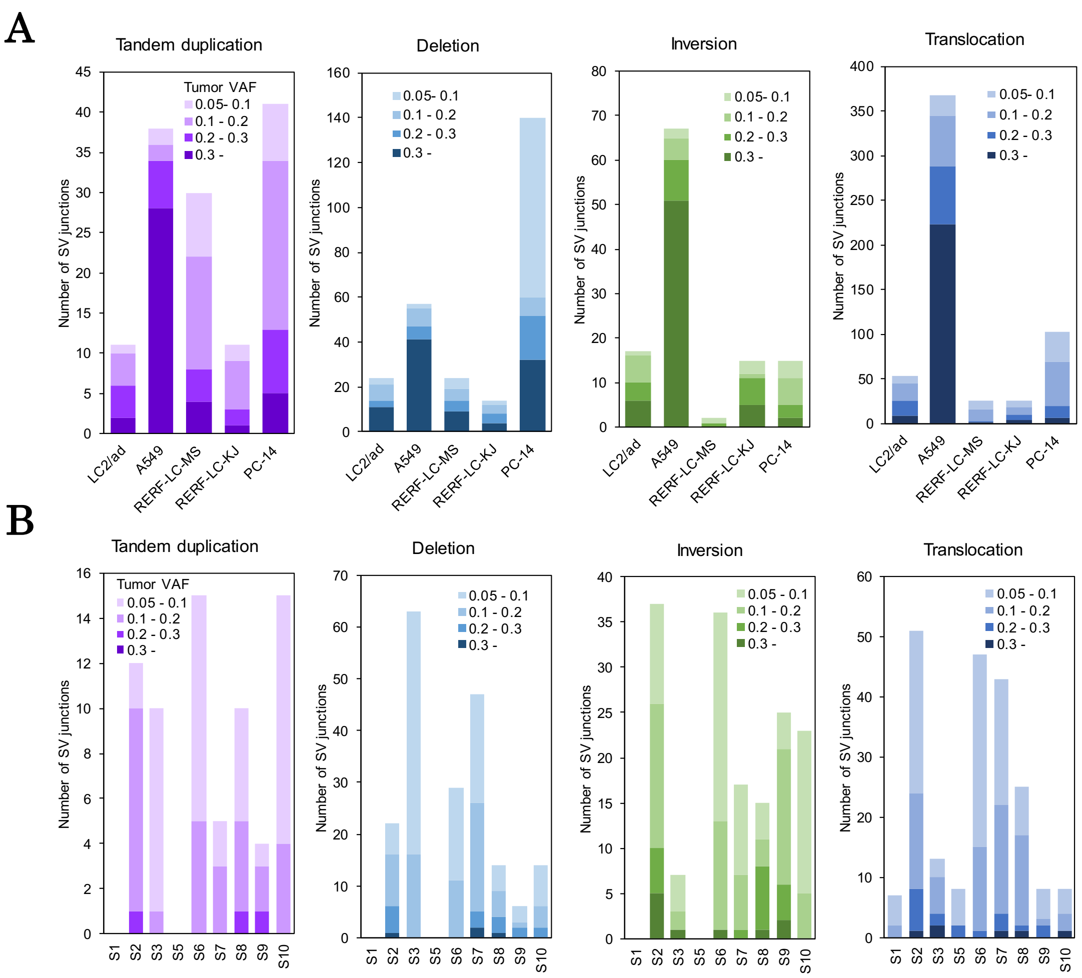


**Supplementary Figure S6 SV junctions detected by GenomonSV.**

(**A B**) SV junctions of the cell lines (**A**) and clinical samples (**B**) were detected by GenomonSV. The colors were graduated according to the tumor variant allele frequencies of each SV.

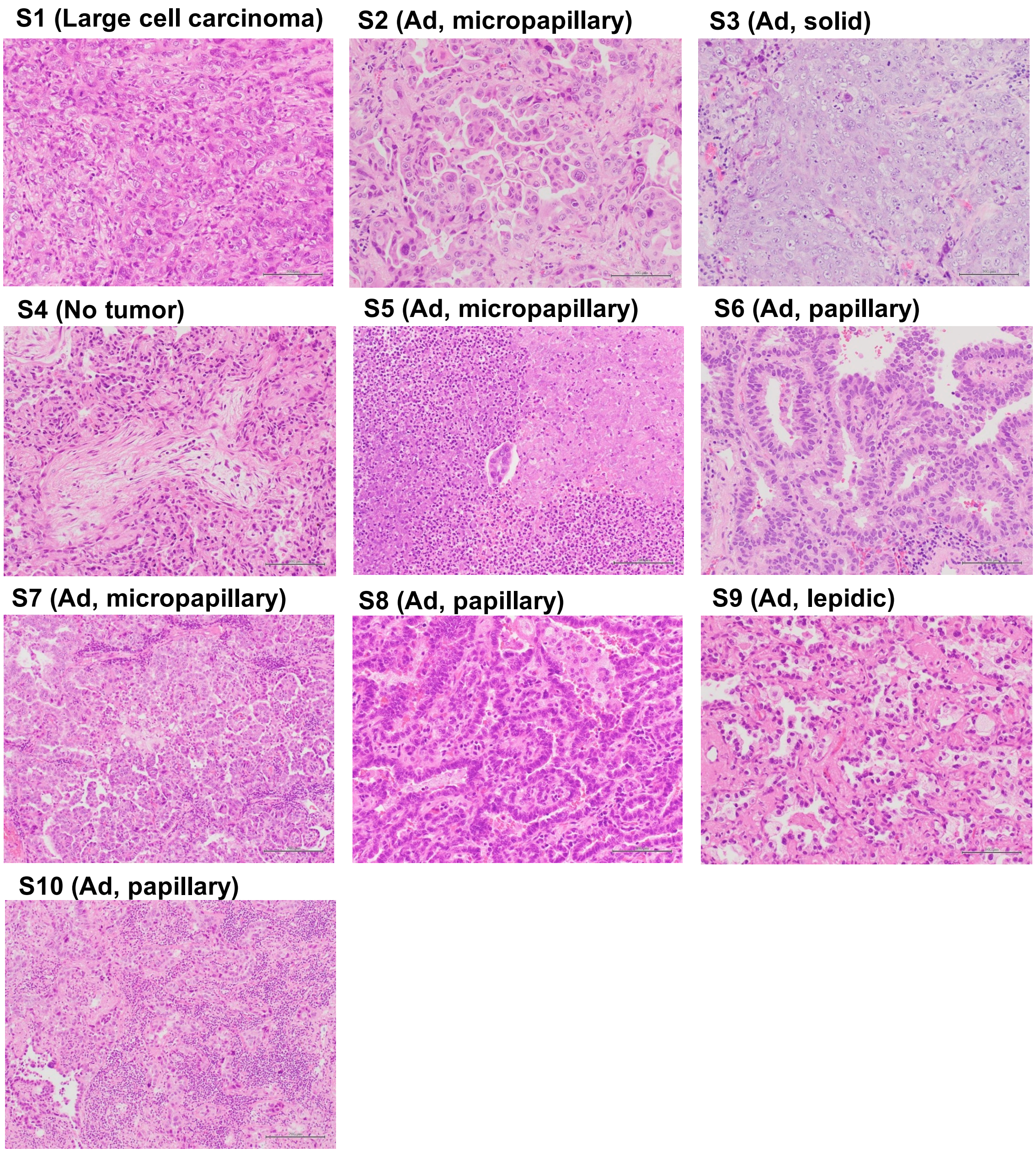


**Supplementary Figure S7 Representative histological images of clinical lung cancer specimens.**

All but one frozen specimen showed the same histology as the final diagnosis: large cell carcinoma (S1) and adenocarcinoma with various subtypes (micropapillary in S2, S5, S7, solid in S3, papillary in S6, S8, S10 and lepidic in S9). Histological examination of Case S4 revealed an organizing pneumonia area near the tumor that resembled the tumor by macroscopic evaluation.

**Supplementary Table S1 Summary of lung cancer cell lines**

| Cell lines | Driver mutation | Ethnicity | Gender |
|---|---|---|---|
| LC2/ad | *CCDC6-RET* fusion | Japanese | Female |
| A549 | *KRAS* G12S | Non-Asian | Male |
| PC-14 | NRAS Q61K | Japanese | Male |
| RERF-LC-MS | Unknown | Japanese | Male |
| RERF-LC-KJ | Unknown | Japanese | Male |

**Supplementary Table S2 Sequencing statistics of MinION and PromethION**

**A. A549**

| Categories | Number |
|---|---|
| Number of reads | 2,059,786 |
| N50 length of reads | 29,535 bp |
| Coverage | 10.1 × |
| Percentage of mapped reads | 80.9 % |
| Average length of mapped reads | 16,888.1 bp |
| Average identity of mapped reads | 80.9 % |

**B. RERF-LC-KJ**

| Categories | Number |
|---|---|
| Number of reads | 5,986,875 |
| N50 length of reads | 23,442 bp |
| Coverage | 18.5 × |
| Percentage of mapped reads | 74.5 % |
| Average length of mapped reads | 12,293.6 bp |
| Average identity of mapped reads | 86.5 % |

**C. RERF-LC-MS**

| Categories | Number |
|---|---|
| Number of reads | 2,261,625 |
| N50 length of reads | 36,418 bp |
| Coverage | 9.5 × |
| Percentage of mapped reads | 74.7 % |

| | |
|---|---|
| Average length of mapped reads | 16,454.4 bp |
| Average identity of mapped reads | 83.0 % |

**D. PC-14**

| Categories | Number |
|---|---|
| Number of reads | 5,967,298 |
| N50 length of reads | 17,804 bp |
| Coverage | 11.5 × |
| Percentage of mapped reads | 74.3 % |
| Average length of mapped reads | 7,698.4 bp |
| Average identity of mapped reads | 85.2 % |

A549: Sequenced by MinION; the others: Sequenced by PromethION

**Supplementary Table S3 Candidate novel fusion genes**

**A. LC2/ad**

| Chr1 | Position1 | Gene1 | Chr2 | Position2 | Gene2 | Nanopore reads | Illumina PE | Illumina soft clipping |
|---|---|---|---|---|---|---|---|---|
| 11 | 21,220,687 | *NELL1* | 4 | 90,671,883 | *CCSER1* | 14 (6, 8) | 13 | 7, 10 |
| 5 | 107,380,583 | *EFNA5* | 8 | 42,299,784 | *IKBKB* | 16 (10, 6) | 6 | 2, 7 |

**B. RERF-LC-KJ**

| Chr1 | Position1 | Gene1 | Chr2 | Position2 | Gene2 | Nanopore reads | Illumina PE | Illumina soft clipping |
|---|---|---|---|---|---|---|---|---|
| 3 | 191,278,568 | *UTS2B* | 6 | 34,029,376 | *GRM4* | 9 | 2 | 4, 6 |

Nanopore reads: the number of nanopore reads supporting the junctions. In LC2/ad cells, the number of both MinION and PromethION reads are also shown (in parentheses: left, MinION; right, PromethION). Illumina PE: the number of Illumina short reads supporting the structures by paired-end data. Illumina soft clipping: the number of Illumina short reads supporting the junctions by soft clipping.

**Supplementary Table S4 Histopathological information on lung cancer clinical samples**

| Case | T/N ratio (tumor purity) | Necrosis | Inflammation |
|---|---|---|---|
| S1 | Moderate | Present | Mild |
| S2 | Moderate | Absent | Moderate |
| S3 | Moderate | Present | Moderate |
| S4* | No tumor | Present | Moderate |
| S5 | Low | Present | Severe |
| S6 | Moderate | Absent | Moderate |
| S7 | Moderate | Absent | Moderate |
| S8 | High | Absent | Moderate |
| S9 | Moderate | Absent | Mild |
| S10 | Low | Absent | Severe |

*Case S4 was removed from the analysis because of low tumor purity.

**Supplementary Table S5 General statistics of PromethION in lung cancer clinical samples**

| Case | Number of Reads | N50 Length of Reads (bp) | Coverage (×) | Percentage of Mapped Reads (%) | Average Length of Mapped Reads (bp) | Average Identity of Mapped Reads (%) |
|---|---|---|---|---|---|---|
| S1 | 5,372,418 | 22,190 | 13.6 | 79.3 | 9,243 | 82.6 |
| S2 | 13,582,749 | 13,565 | 16.6 | 81.3 | 4,419 | 86.5 |
| S3 | 7,506,147 | 15,977 | 12.1 | 72.6 | 6,467 | 86.2 |
| S5 | 10,275,907 | 12,496 | 13.4 | 75.7 | 5,018 | 86.0 |
| S6 | 6,387,904 | 20,731 | 14.0 | 71.5 | 8,912 | 86.7 |
| S7 | 7,107,011 | 17,478 | 13.7 | 72.0 | 7,706 | 86.0 |
| S8 | 6,457,028 | 17,938 | 11.1 | 73.0 | 6,797 | 86.5 |
| S9 | 7,855,858 | 21,101 | 18.1 | 76.1 | 8,711 | 86.4 |
| S10 | 6,423,471 | 21,433 | 15.6 | 78.0 | 8,902 | 84.2 |

**Supplementary Table S6 Numbers of genes affected by CLCLs in cell lines and clinical samples**

**A. Cell lines**

| Cell | Number of genes with CLCLs | | |
|---|---|---|---|
| | TP (both detected) | FP (short read only) | FN (long read only) |
| LC2/ad | 11 | 2 | 6 |
| A549 | 2 | 33 | 0 |
| RERF-LC-MS | 6 | 24 | 1 |
| RERF-LC-KJ | 3 | 9 | 5 |
| PC-14 | 11 | 29 | 1 |
| Total | 33 | 97 | 13 |

**B. Clinical samples**

| Case | Number of genes with CLCLs | | |
|---|---|---|---|
| | TP (both detected) | FP (short read only) | FN (long read only) |
| S1 | 0 | 0 | 0 |
| S2 | 2 | 10 | 1 |
| S3 | 1 | 11 | 0 |
| S5 | 0 | 0 | 0 |
| S6 | 0 | 14 | 1 |
| S7 | 2 | 4 | 0 |
| S8 | 2 | 9 | 0 |
| S9 | 1 | 3 | 0 |
| S10 | 3 | 17 | 2 |
| Total | 11 | 68 | 4 |

TP: true positives detected in both long read and short read data; FP: false positives detected in short read data only; FN: false negatives detected in long read data only.
